# Audiovisual task switching rapidly modulates sound encoding in mouse auditory cortex

**DOI:** 10.1101/2021.11.09.467944

**Authors:** Ryan J. Morrill, James Bigelow, Jefferson DeKloe, Andrea R. Hasenstaub

## Abstract

In everyday behavior, sensory systems are in constant competition for attentional resources, but the cellular and circuit-level mechanisms of modality-selective attention remain largely uninvestigated. We conducted translaminar recordings in mouse auditory cortex (AC) during an audiovisual (AV) attention shifting task. Attending to sound elements in an AV stream reduced both pre-stimulus and stimulus-evoked spiking activity, primarily in deep layer neurons. Despite reduced spiking, stimulus decoder accuracy was preserved, suggesting improved sound encoding efficiency. Similarly, task-irrelevant probe stimuli during intertrial intervals evoked fewer spikes without impairing stimulus encoding, indicating that these attention influences generalized beyond training stimuli. Importantly, these spiking reductions predicted trial-to-trial behavioral accuracy during auditory attention, but not visual attention. Together, these findings suggest auditory attention facilitates sound discrimination by filtering sound-irrelevant spiking in AC, and that the deepest cortical layers may serve as a hub for integrating extramodal contextual information.

## Introduction

Information from one or another sensory pathway may become differentially relevant due to environmental changes. The brain must therefore continuously assign limited attentional resources to processing simultaneous information streams from each sensory modality. For example, hearing a siren while listening to music in the car might prompt an attention shift away from the auditory stream toward a visual search for emergency vehicles. On the other hand, a similar shift away from the music is unlikely while listening at home. In these cases, contextual cues support allocating attention to either the auditory domain or the visual domain, and the perceptual experience of the music is qualitatively different. How might sensory cortex differentially encode stimuli from an attended versus filtered modality?

Attentional selection operates cooperatively at many levels of sensory processing. Most effort has been devoted to understanding the neural mechanisms of feature-selective attention within a single modality (Desimone and Duncan, 1995; Fritz et al., 2007). A major focus of this work has been characterizing transformations of stimulus representations in sensory cortical areas, due to their pivotal position between ascending sensory pathways and behavioral networks implementing top-down control (Lamme et al., 1998; Sutter and Shamma, 2011). These studies, largely from the visual domain, have shown that attention to a stimulus feature or space will often increase stimulus-evoked spiking responses and reduce thresholds for eliciting a response; likewise, responses to unattended stimuli are often decreased (Reynolds and Chelazzi, 2004). On the other hand, fewer studies have examined how modality-selective attention affects encoding in sensory cortex. This mode of attention highlights behaviorally-relevant sensory streams while filtering less relevant ones. Human fMRI studies have reported differential activation patterns in auditory and visual cortex (AC, VC) reflecting the attended modality (Johnson and Zatorre, 2005; Petkov et al., 2004; Shomstein and Yantis, 2004; Woodruff et al., 1996). Extending these findings, studies in primate AC and VC have reported entrainment local field potential (LFP) oscillations by modality-selective attention, which serves to modulate excitability and sharpen feature tuning within sensory cortex corresponding to the attended modality (Hocherman et al., 1976; Lakatos et al., 2008, 2009; O’Connell et al., 2014). Several findings suggest these influences may differ among cortical layers and between inhibitory and excitatory neurons (Lakatos et al., 2016; O’Connell et al., 2014).

Nevertheless, much remains unknown about the influence of modality-specific attention on stimulus encoding in sensory cortex. Importantly, potential interplay between ongoing activity and evoked responses, as well as their consequences for information and encoding efficiency has not been examined. How these influences may be differentially expressed in cell subpopulations defined by cortical depth or inhibitory/excitatory cell type similarly remains unknown. Finally, the degree to which influences of modality-specific attention may generalize beyond training stimuli remains unknown.

In the present study, we addressed these open questions by examining single neuron activity and sensory responses in mouse AC during an audiovisual attention shifting task. AC integrates ascending auditory information with diverse input from frontal, cingulate, striatal, and non-auditory sensory areas to rapidly alter sensory processing in response to changing behavioral demands (Budinger and Scheich, 2009; Budinger et al., 2008; Park et al., 2015; Rodgers and DeWeese, 2014; Winkowski et al., 2013). To isolate the influence of modality-selective attentional modulation, we compared responses to identical compound auditory-visual stimuli within task, under different cued contexts requiring attention to the auditory or visual elements, thus holding constant other task-related variables such as arousal, attention, reward expectation, and motor activity (Saderi et al., 2021). Because spike rate and information changes are dissociable (Bigelow et al., 2019; Phillips and Hasenstaub, 2016), we quantified both evoked rates and mutual information between responses and stimuli. We also examined the generality of modality-specific attention by examining responses to task-irrelevant sounds presented between trials. Finally, we used translaminar probes and spike waveform morphology classification to capture possible attention-related differences in neurons among cortical layers and between putative inhibitory and excitatory cell classes.

## Results

### Audiovisual rule-switching in mice

We trained mice to perform an AV rule-switching task, in which they made decisions using auditory stimuli while ignoring simultaneously presented visual stimuli or vice versa. Trial presentation was self-paced in a virtual foraging environment wherein a visual track was advanced by forward locomotion on a spherical treadmill (Fig. 1.A). A task-irrelevant random double sweep (RDS) sound was presented during intertrial intervals (ITIs) for mapping auditory receptive fields in each attentional state (**Fig. 1.B**). Decision stimuli were presented after variable track length, comprising 1-s auditory tone clouds (centered at 8 or 17 kHz) and/or visual drifting gratings (horizontal or vertical orientation). One of the decision stimuli for each modality was a rewarded target (A_R_, V_R_) and the other an unrewarded distractor (A_U_, V_U_). Lick responses following targets (hits) and distractors (false alarms; FAs) produced water rewards and dark timeouts, respectively. Withholding licks for targets (misses) or distractors (correct rejects; CRs) advanced the next trial. Each session began with a block of unimodal decision stimuli, which cued the attended modality of a subsequent AV block (**Fig. 1.D**). A second unimodal block from the other modality was then presented, cueing the rule for a final AV block. Decision stimuli had identical physical properties but different behavioral significance between rules (e.g., licks following A_R_V_U_ were rewarded in A-rule but punished in V-rule). Targets and distractor stimuli remained constant throughout training for each mouse and were approximately counterbalanced across animals. Block sequences (A-rule then V-rule, or vice versa) were also counterbalanced across sessions.

**Fig. 1.**
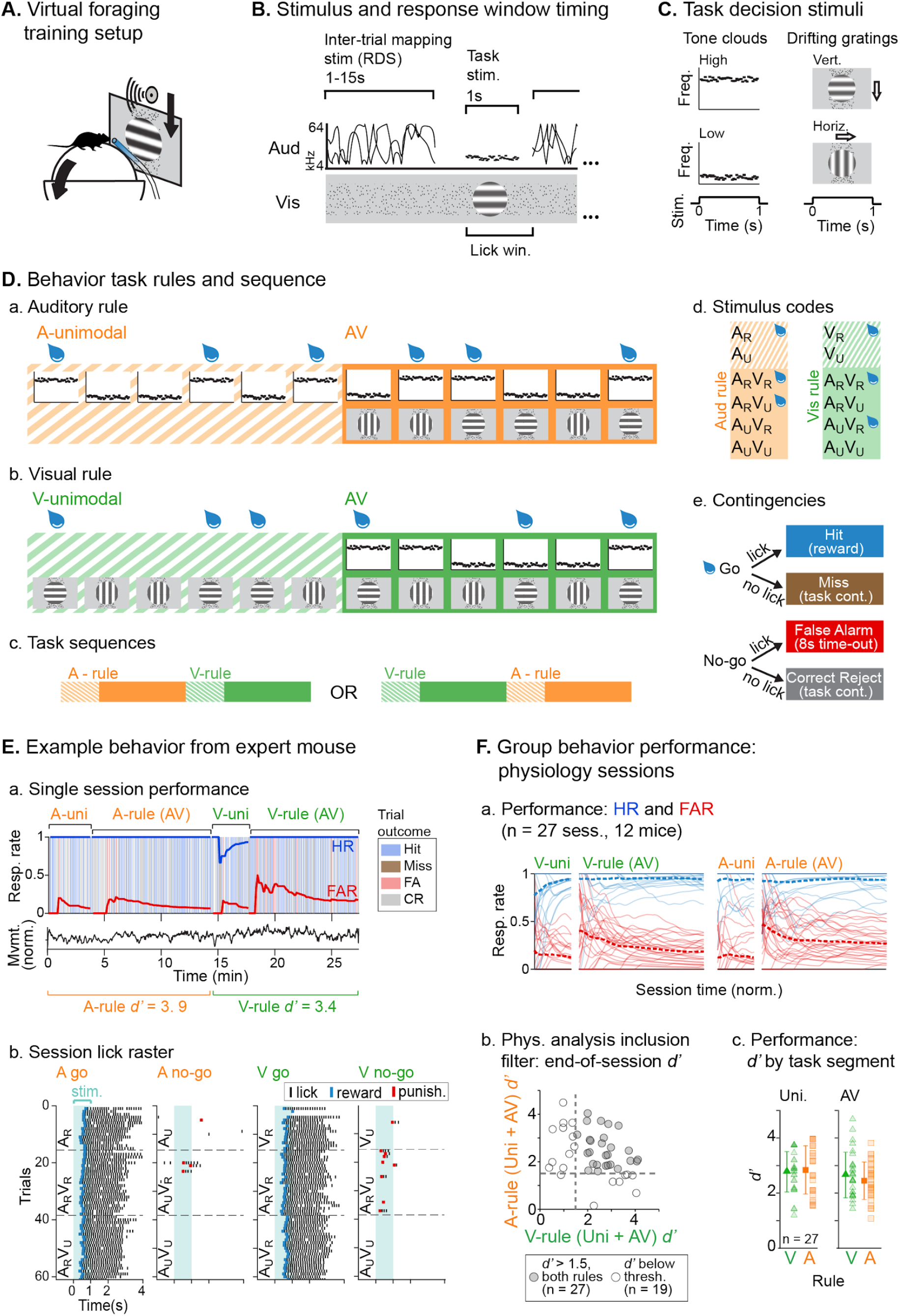
Audiovisual rule-switching in mice. **A.** Virtual foraging environment. Mice advance a track display by forward movement and respond to decision stimuli with lick responses for water rewards. **B.** Trial sequence including decision stimuli and intertrial intervals featuring visual track dots and auditory mapping stimulus. **C.** Decision stimuli comprised 1-s tone clouds (8 or 17 kHz) and/or drifting gratings (horizontal or vertical). **D.** Task sequences, attention cueing, and reward contingencies, **a–b.** A-rule vs. V-rule was cued by a unimodal block presenting auditory or visual decision stimuli alone, **c.** Each session reflected one of two possible task sequences, **d.** Stimulus codes reflecting each possible combination of rewarded (R) and unrewarded (U) stimuli within the auditory (A) and visual (V) modalities, presented in unimodal or AV blocks, **e.** Contingencies for water reward, time-out punishment, or task continuation. **E.** Example behavior session, a. Behavioral outcomes for each trial and attention condition, b. Lick responses by condition and stimulus. **F.** Task performance summary for sessions included in physiology analyses, a. HR and FAR by rule for each session (dashed lines, mean), **b.** Sensitivity index *d’* by rule for all sessions, **c.** Average performance (*d’*, mean ±SD) for sessions with *d’*>1.5 for both rules, which were included in physiology analyses.

We used two approaches to ensure that animals were engaged during both task rules. First, we restricted analysis to sessions in which discrimination was well above chance (*d’* >1.5) for both rules (**Fig. 1.F**). Second, for a subset of sessions (n = 14 sessions, 5 mice) we measured pupil size, a well-established correlate of arousal and behavioral performance (Bradley et al., 2008; McGinley et al., 2015; Reimer et al., 2014). We used a computer vision algorithm to automate pupil size (pupil circumference / eye diameter) measurement for each frame acquired by a CCD video camera (**Fig 2.A.a**). To isolate pupil fluctuations reflecting general arousal, pupil size was measured during an inter-trial interval (ITI) window designed to avoid pupil responses to decision stimulus onset, dark timeouts, and decreased locomotion following reward administration (**Fig. 2.A.b-c, Fig 2.B**). Consistent with previous studies reporting pupil size increases with task difficulty (Hess and Polt, 1964; Kawaguchi et al., 2018), pupil size was significantly larger for bimodal compared to unimodal blocks within each rule (**Fig. 2.C**; A-rule unimodal: 0.284 ±0.041 [mean ±SD], A-rule bimodal: 0.293 ±0.047; Z = −2.574, p = 0.005, paired Wilcoxon signed rank [WSR], one-tailed; V-rule unimodal: 0.292 ±0.044, V-rule bimodal: 0.297 ±0.045; Z = −2.009, p = 0.022). By contrast, no difference in pupil size was observed between rules during bimodal blocks (**Fig. 2.C**; A-rule bimodal: 0.293 ±0.047, V-rule bimodal: 0.297 ±0.045; Z = −1.036, p = 0.30, paired WSR, two-tailed). Because pupil size closely tracks locomotion (**Fig. 2.A.b-c**; McGinley et al., 2015), we also examined locomotion speed during the same ITI window (**Fig 2.D**). Differences in locomotion speed were neither observed between unimodal and bimodal blocks within rule nor between bimodal blocks between rules (all p ≥ 0.623; all |z| ≤ 1.29, paired WSR). Arousal and motor activity were thus comparable between rules, isolating any differences in neuronal activity to modality-selective attention.

**Fig. 2.**
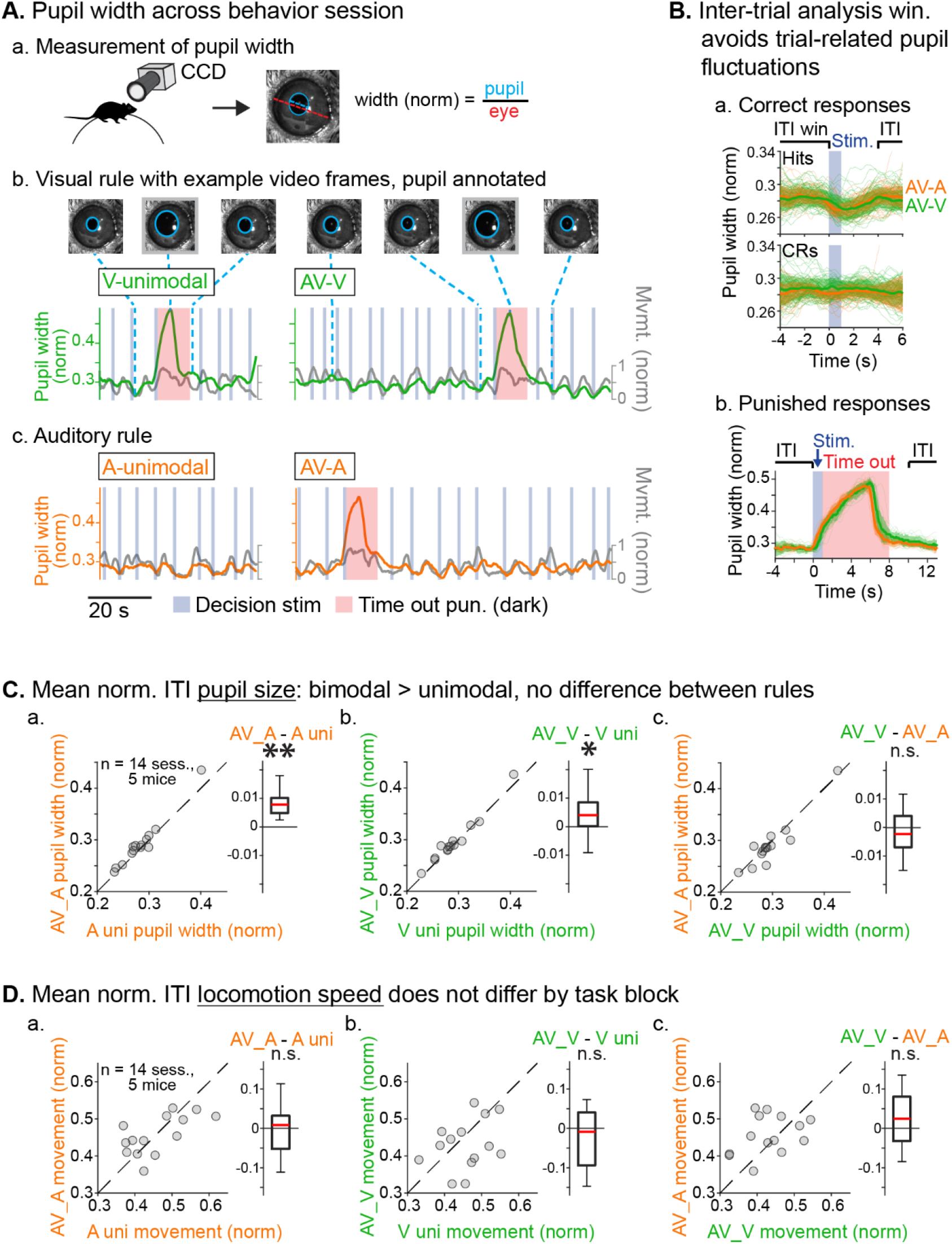
Similar levels of arousal and movement during auditory and visual attention. **A.** Pupil size measurement, **a.** Pupil recorded via CCD camera; size is measured as pupil diameter by visible eye diameter, **b–c.** Example pupil video frames showing correlated pupil size and movement measurements during example task trials. Large pupil size increases were typical of dark timeout punishments. **B.** Pupil size during ITIs serve as an index of arousal, **a–b.** Pupil size decreases and increases typically followed hits (reward delivery) and FAs (timeouts), respectively. The ITI analysis window avoids these influences. **C.** Mean pupil size by condition, **a–b.** Pupil size was larger during AV than unimodal blocks, suggesting increased arousal during more difficult task segments, c. Pupil size did not differ between attention conditions (AV blocks), suggesting similar levels of arousal. **D.** Mean movement velocity by condition, **a–c.** Movement did not differ between unimodal and AV blocks or between attention conditions, suggesting comparable locomotor activity across conditions and indicating pupil increase during AV blocks were not driven by increased movement.

### Single unit recording in AC

After mice learned the AV rule-switching task, a craniotomy was made over right AC for acute recordings during behavior using multichannel probes spanning the full cortical depth (**Fig. 3.A**). In total, we recorded AC activity in 12 mice during 27 behavioral sessions meeting inclusion criteria. The putative cortical depth of each sorted single unit (SU) was assigned by calculating the fractional position of the channel with the largest waveform amplitude within the span of channels in AC as estimated from spontaneous and tone-evoked recordings following the task (**Fig. 3.B**). A separate set of experiments using probe track visualization with Di-I provided support for this depth estimation technique (**Fig. 3.C**; DiCarlo et al., 1996; Morrill and Hasenstaub, 2018). We then divided the fractional depth values into superficial, middle, and deep groups, approximating the supragranular, granular, and infragranular laminae. We further divided SUs into narrow-spiking (NS, putative inhibitory; n = 150, 17%) and broad-spiking (BS, predominantly excitatory; 723, 83%) populations based on trough-peak time (**Fig. 3.D**; Cardin et al., 2007; Nandy et al., 2017; Phillips et al., 2017; Bigelow et al., 2019).

**Fig. 3.**
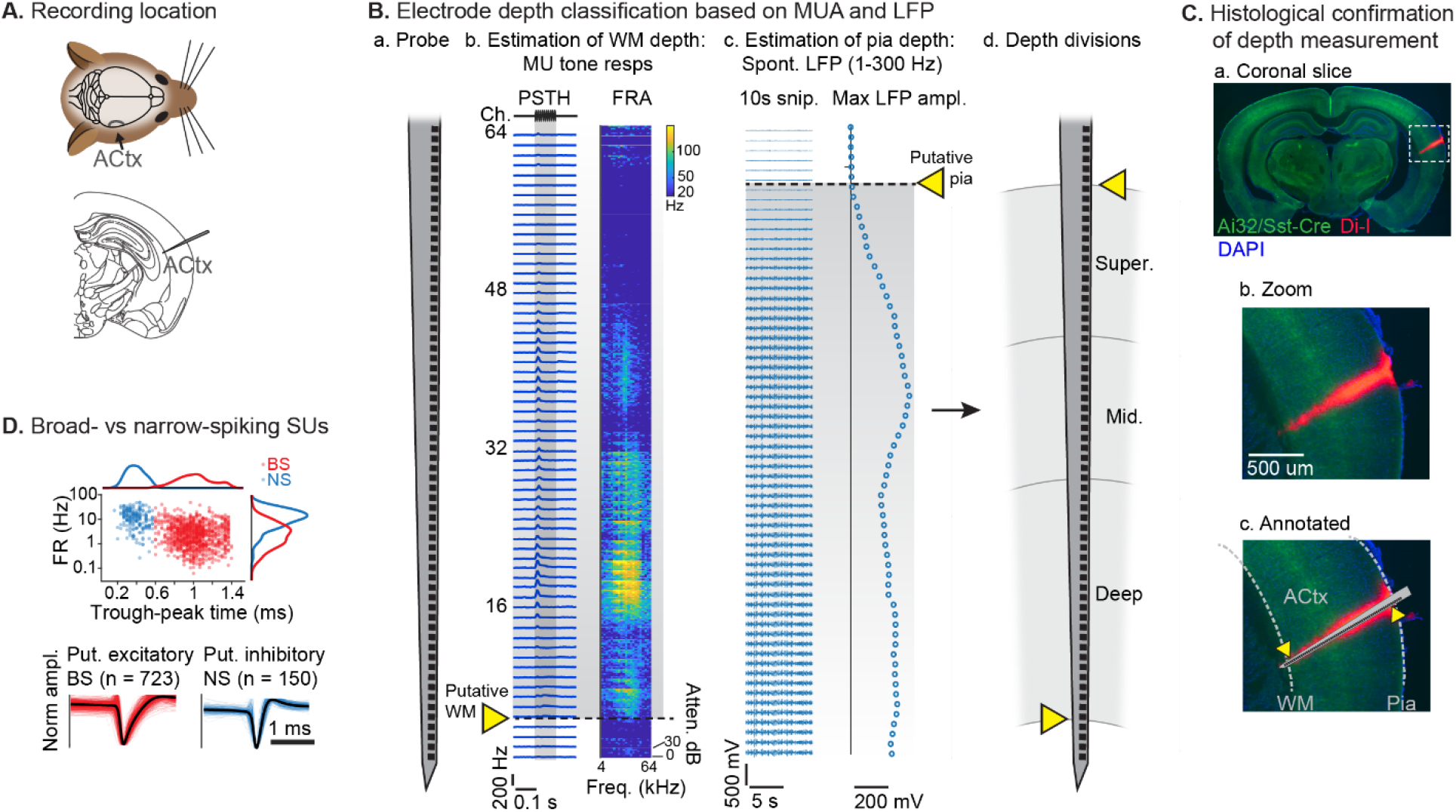
Single unit recording and depth estimation in auditory cortex. **A.** Translaminar probes recorded activity in right AC. **B.** Physiological estimation of cortical depth, **a.** Linear 64-channel probe used to record across AC layers, **b.** Example tone-evoked multi-unit (MU) responses by channel. Left: PSTH plots showing mean responses by time. Right: frequency response area (FRA) plots showing mean responses by tone frequency/attenuation. MU responses provided a marker for the lower cortex-white matter (WM) boundary, but often poorly estimated the upper cortex-pia boundary, likely reflecting minimal somatic spiking in the superficial layer, **c.** Local field potential (LFP; 1-300 Hz) amplitude provides a more reliable a marker for the upper cortex-pia boundary. Left: Example LFP activity by channel. Right: maximum LFP amplitude by channel. Upper cortex-pia boundary was estimated by the first meaningful deviation from probe-wise minimum LFP amplitude, d. Relative cortical depth reflected fractional division of cortex into ‘superficial’, ‘middle’, and ‘deep’, with fractions based on supragranular, granular and infragranular anatomical divisions. **C.** Histological confirmation of cortical depth estimation technique, **a.** Coronal slice showing Di-I probe marking in right AC. Green: eYFP fluorescence in Ai32/Sst-Cre mouse; Blue: DAPI cell body staining, b. Zoomed area indicated by dashed rectangle in a. c. Probe overlay with WM and pia boundaries. **D.** Sorted SU waveforms were divided into narrow-spiking (putative fast-spiking inhibitory) and broad-spiking (putative excitatory) populations based on a waveform trough-peak time boundary of 0.6 ms.

### Modality-selective attention modulates stimulus-evoked firing rates

Our analysis of SU activity during AV rule-switching focused on comparisons of responses to bimodal decision stimuli between rules, which reflected physically identical stimuli and comparable arousal and locomotion levels. Responses evoked by decision stimuli and modulatory effects of task rule were diverse (Fig. 4.A). To capture a predominantly sensory-driven component of the response, we measured firing rates (FRs) within the first 300 ms post-stimulus onset (**Fig. 4.B**), which preceded most lick responses (lick latency median: 583 ms; 5th-95th percentiles: 275-1064 ms; 5.8% of licks <300 ms, n = 3,342 total lick trials across dataset). Trials with licking activity earlier than 300 ms were excluded from analysis. We first compared A-rule and V-rule responses to stimuli rewarded in the A-rule (A_R_*: A_R_V_R_ and A_R_V_U_ responses combined). Averaging across units, responses in the middle and deep layers were suppressed in the A-rule relative to the V-rule for both NS and BS units (**Fig. 4.C** Mid. BS: p = 0.011, Z = 2.55, median fold change [FC; A-rule/V-rule]: 0.87, n = 153 SUs; Mid. NS: p = 0.0096, Z = 2.59, med. FC: 0.80, n = 34; Deep BS: p = 2.7e-06, Z = 4.70, med. FC: 0.88, n = 391; Deep NS: p = 0.0015, Z = 3.17, med. FC:0.87, n = 77; paired Wilcoxon signed rank [WSR]; see **Table S1.A** for full stats). No significant group-level change was found in superficial units. Consistent with group level trends, individual units with significant FR decreases in the A-rule (p < 0.01, unpaired t-test) outnumbered units with significant FR increases for all unit populations other than superficial and mid-depth BS units (**Fig. 4.C**, right).

**Fig. 4.**
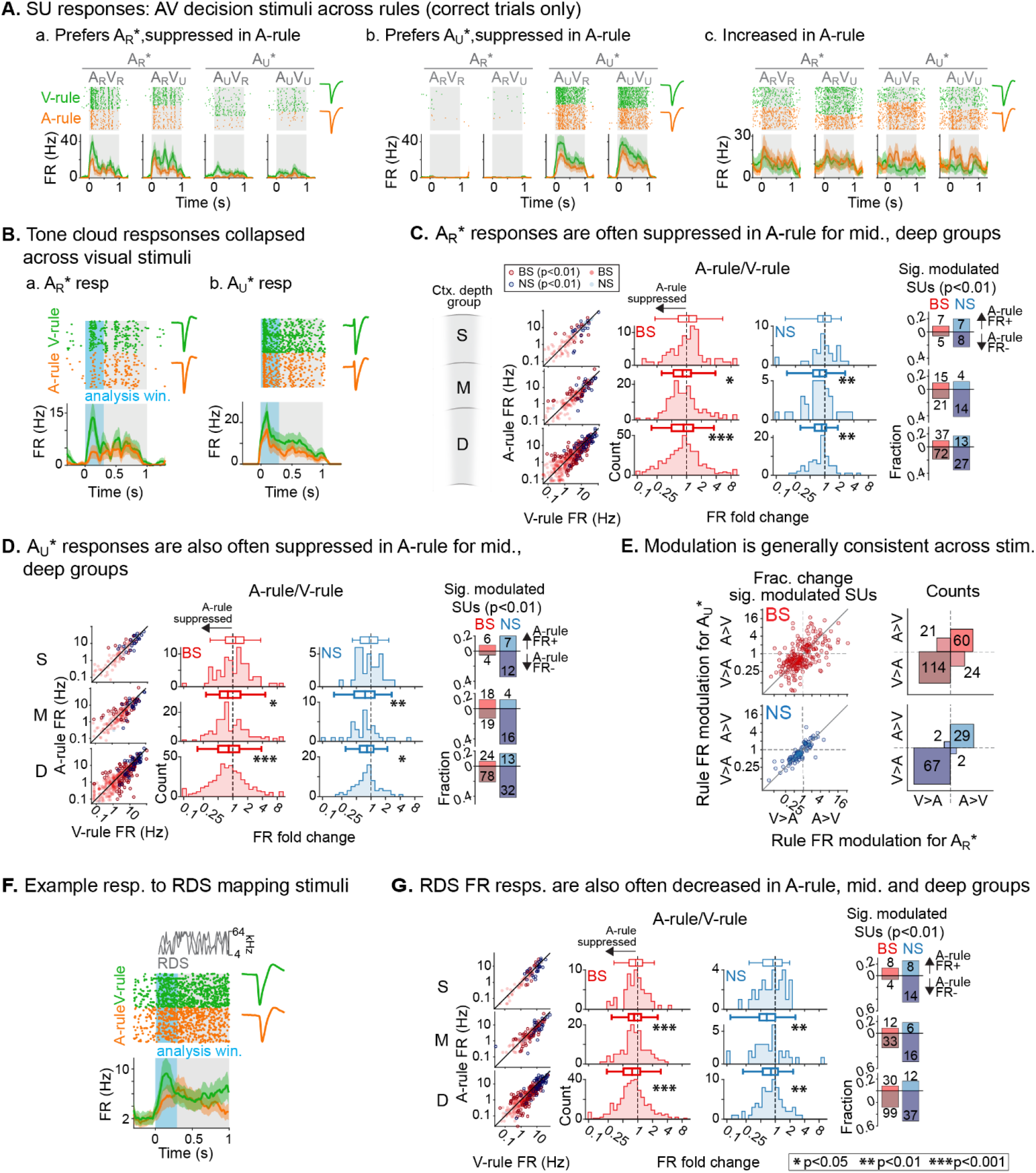
Net suppression of sound-evoked firing rates during auditory attention. **A.** Example units with attention-modulated responses to sound. (A_R_*: A_R_V_R_ and A_R_V_U_ collapsed; A_U_*: A_U_V_R_ and A_U_V_U_ collapsed). **B.** Example SU responses to rewarded and unrewarded sounds between rules collapsed across visual stimuli. Early sensory response analysis window shown in blue (0-0.3 s). **a.** Rewarded sound (A_R_*: A_R_V_R_ and A_R_V_U_ collapsed), **b.** Unrewarded sound (A_U_*: A_U_V_R_ and A_U_V_U_ collapsed). **C.** Group data: responses to rewarded sounds (A_R_*) between rules by unit type and depth. Left: Individual unit responses by rule (outline: significantly modulated, p < 0.01; paired t-test). Middle: Histograms of FR fold change between rules (A-rule / V-rule); values <1 indicate A-rule suppression. Box plots indicate median, 25th-75th percentiles, and range excluding outliers. Right: Fractions of significantly modulated units; values above and below the zero line reflect increased and decreased FRs in the A-rule, respectively. **D.** Group data: responses to rewarded sounds (A_U_*) between rules (conventions as in C). **E.** Comparison of unit FR modulation between A_R_* (abscissa) and A_U_* (ordinate). Left: Modulation by stimulus for individual units with significant modulation for one or both sounds; values <1 indicate A-rule suppression. Right: counts of units with significantly increased and decreased FRs by stimulus. Most units showed similar modulation direction across stimuli. **F.** Example SU responses to task-irrelevant sounds during the ITI between rules. **G.** Group data: responses to task-irrelevant sounds between rules (conventions as in C).

A strikingly similar pattern of attention-related modulation was observed for unrewarded stimuli in the A-rule (A_U_*: A_U_V_R_ and A_U_V_U_ responses combined). At the group level, superficial unit responses did not significantly differ between conditions, whereas middle and deep NS and BS units were significantly suppressed in the A-rule (**Fig. 4.C.b**; Mid. BS: p = 0.015, Z = 2.42, med. FC: 0.81, n = 148; Mid. NS: p = 0.0025, Z = 3.02, med. FC: 0.76, n = 34; Deep BS: p = 2.7e-08, Z = 5.56, med. FC: 0.82, n = 374; Deep NS: p = 0.017, Z = 2.38, med. FC: 0.83, n = 77; paired WSR; see **Table S1.B** for full stats). Relative fractions of units with significantly increased and decreased FRs to A_U_* stimuli were similar to those described above for A_**R**_* stimuli (**Fig. 4.D**, right). We further found that most units showed the same direction of modulation for A_R_* and A_U_* stimuli (**Fig. 4.E**), with similar modulation sign observed for 79% of BS units (52% suppressed for both A_R_* and A_U_*, 27% enhanced for both) and 96% of NS units (67% suppressed for both, 29% enhanced for both). These findings suggest modality-selective attention similarly influences FRs evoked by task-relevant target and distractor sounds with different acoustic properties and learned behavioral values.

To determine whether these attentional influences might generalize to task-irrelevant sounds, we examined responses to RDS sounds presented during the ITI. Using the same analysis window (300 ms post-stimulus onset, Fig. 4.F), we found that attention-related modulation of FR responses evoked by task-irrelevant sounds was highly similar to that observed for both types of decision stimuli: middle- and deep-layer BS and NS populations exhibited group-level FR suppression during the A-rule (Fig. 4.G), whereas superficial layer units were not significantly modulated (Mid. BS: p = 9e-05, Z = 3.92, med. FC: 0.86, n = 134; Mid. NS: p = 0.0037, Z = 2.90, med. FC: 0.68, n = 33; Deep BS: p = 3.2e-12, Z = 6.97, med. FC: 0.81, n = 350; Deep NS: p = 0.0079, Z = 2.65, med. FC: 0.83, n = 72; paired WSR; see Table S2 for full stats). The majority of individual units with significant FR changes reflected A-rule suppression for all depth and unit type subpopulations other than superficial BS units. The difference was again most pronounced for deep layer units (BS: 8% sig. enhanced, 28% suppressed; NS: 15% enhanced, 52% suppressed). Together, these results show that auditory-selective attention tends to reduce FR responses to sounds, regardless of their behavioral relevance, valence, or spectral content, and that these influences are strongest for deep layer units.

### Modality-selective attention also modulates pre-stimulus firing rates

Previous studies have found that modulation of ongoing activity in sensory cortex can influence subsequent sensory-evoked responses (Arieli et al., 1996; Haider and McCormick, 2009). Thus, the response suppression during auditory attention reported above may either reflect specific decreases in stimulus responsivity or general decreases in ongoing activity. To address these possibilities, we quantified FRs in a pre-stimulus window spanning 300 ms prior to decision stimulus onset in which no sounds were presented (**Fig. 5.A.a**). Although this window may include anticipatory modulation of activity (Egner et al., 2010; Samuelsen et al., 2012; Cox et al., 2019), it nevertheless provides a measure of baseline activity for comparison with evoked responses. Similar to sound-evoked FR changes, we observed significant group-level decreases in pre-stimulus FRs during the A-rule for BS and NS units in the middle- and deep-layers, but no modulation of superficial units (**Fig. 5.B**; Mid. BS: p = 0.0014, Z = 3.19, med. FC: 0.88, n = 154; Mid. NS: p = 0.0055, Z = 2.78, med. FC: 0.76, n = 34; Deep BS: p = 4.4e-08, Z = 5.47, med. FC: 0.86, n = 403; Deep NS: p = 0.02, Z = 2.32, med. FC: 0.86, n = 77; paired WSR; see **Table S3** for full stats). To test whether the reduction in pre-stimulus FR was sufficient to account for stimulus-evoked changes reported above, we recalculated FRs evoked by decision stimuli as fold change from pre-stimulus FRs (**Fig. 5.A.b**). Following this adjustment, the middle- and deep-layer unit population responses no longer differed between rules (**Fig. 5.C**). Interestingly, after adjusting for pre-stimulus rate we observed modest A-rule suppression in the superficial NS population, with median FR reductions of 16% for A_R_* (p = 0.015, Z = 2.42, n = 35; paired WSR; for other groups see **Table S4.A**) and 12% for A_U_* (p = 0.0059, Z = 2.75, n = 35; paired WSR; see **Table S4.B**). Together, these results suggest that group level decreases in evoked FRs during A-rule are largely due to generalized suppression of ongoing AC activity.

**Fig. 5.**
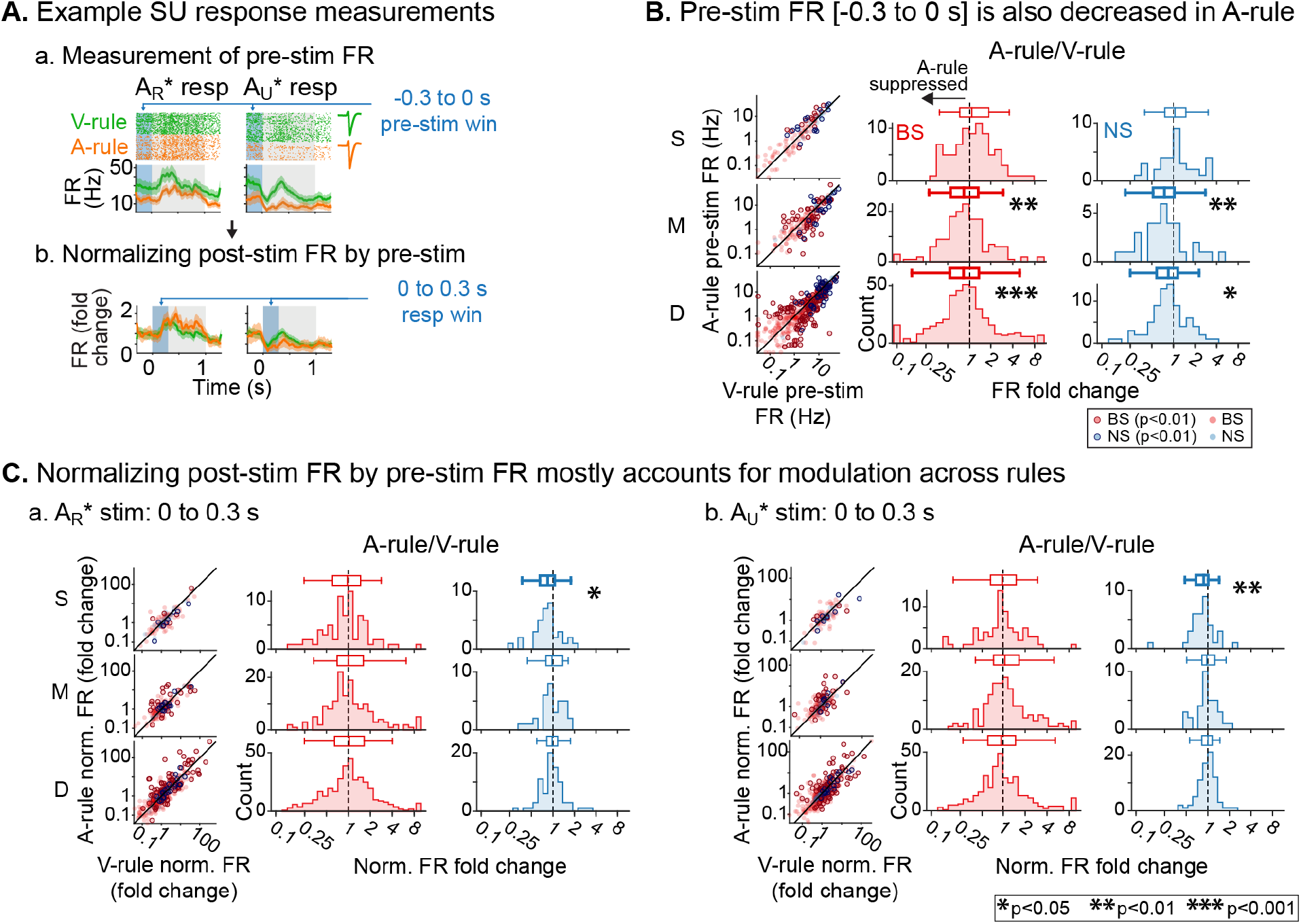
Attention-related modulation of sound-evoked responses largely reflects pre-stimulus activity. **A.** Example unit with attention-modulated pre-stimulus FR. **a.** Raw FRs by condition and stimulus. Pre-stimulus FR analysis window shown in blue (−0.3 to 0 s). b. Pre-stimulus normalized FRs (FR divided by pre-stimulus FR). **B.** Group data: pre-stimulus FRs between rules by unit type and depth (conventions as in Fig. 4C). **C.** Group data: responses to sounds normalized by pre-stimulus FR between rules (conventions as in B). **a.** Responses to rewarded sound (A_R_*). b. Responses to unrewarded sound (A_U_*).

### Attention to sound increases encoding efficiency in deep-layer BS units

Previous work has established that FR changes do not necessarily imply changes in stimulus-spike information. For instance, optogenetic activation of inhibitory interneurons can reduce FRs in AC without changing information, suggesting increased encoding efficiency (Phillips and Hasenstaub, 2016). By contrast, locomotion reduces both FRs and information in AC (Bigelow et al., 2019). To determine whether reduced FRs evoked by decision stimuli were accompanied by changes in information and/or encoding efficiency, we used a PSTH-based neural pattern decoder to compare sound discrimination across attentional states (Foffani and Moxon, 2004; Malone et al., 2007; Hoglen et al., 2018). For each unit, the decoder generates a single-trial test PSTH and then compares these to two or more template PSTHs from distinct stimulus response conditions, generated sans test trial (**Fig. 6.A**). The test trial is assigned to the template that is closest in *n*-dimensional Euclidean space, reflecting *n* PSTH bins. This is repeated for all trials, generating new templates for each classifier run. After all trials have been classified, a confusion matrix is generated. From this, we calculated accuracy of classification, mutual information (MI; bits) and encoding efficiency, a spike-rate normalized MI (bits/spike). As in previous analyses, a 0-300ms post-stimulus onset window was used in this method to restrict decoding to a predominantly sensory-driven component of the response. The binwidth for generating PSTHs was 30 ms (Hoglen et al., 2018). Only trials with correct responses (hits and CRs) and units with a minimum stimulus response FR of 5 Hz to at least one stimulus of the decoder comparison were included.

**Fig. 6.**
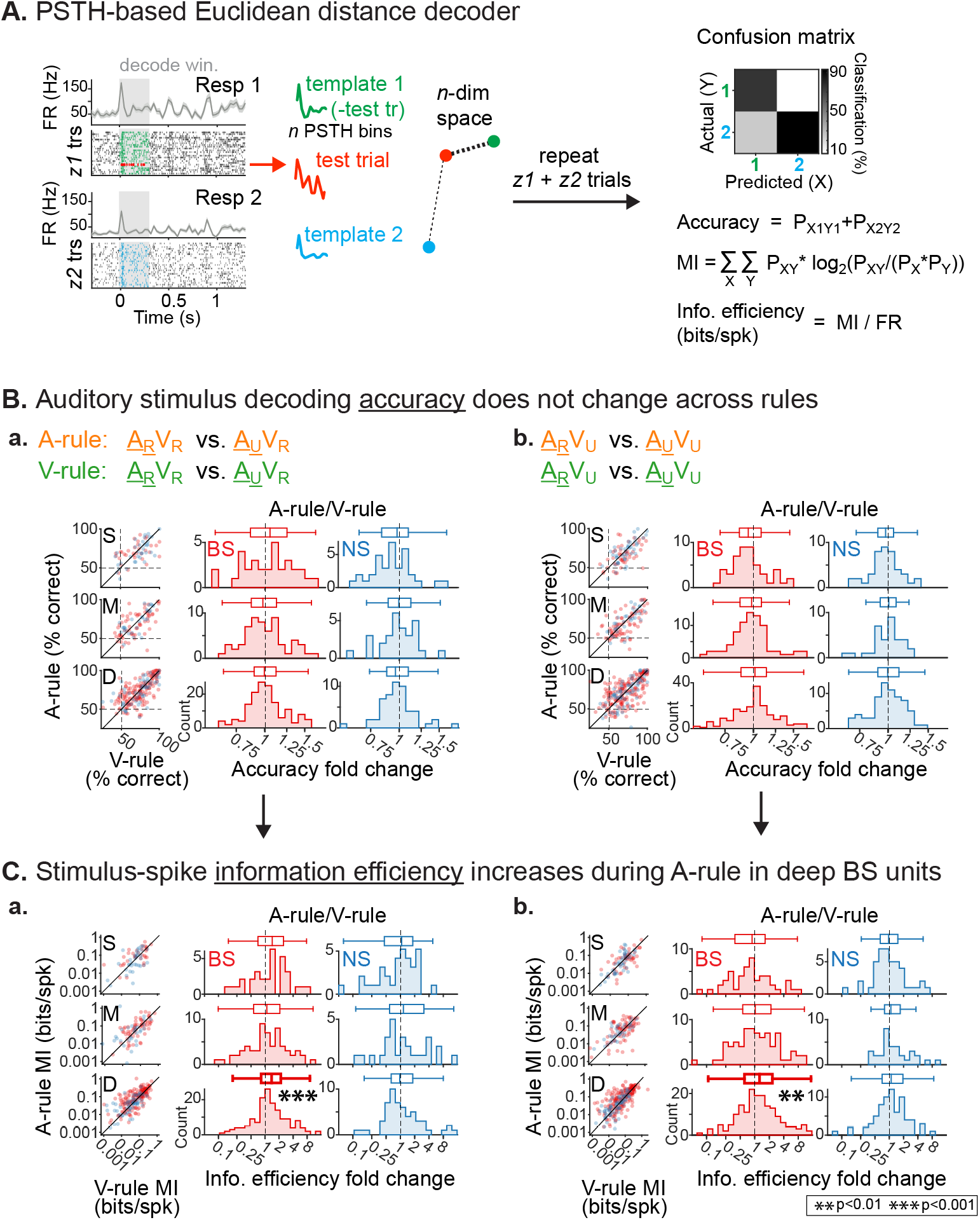
Auditory attention increases encoding efficiency for task-relevant sounds in deep-layer BS units. **A.** Sound classifier analysis. Time-binned responses for each single trial (test trial) are compared to PSTHs (templates) reflecting responses to each sound averaged across all other trials (Resp 1, Resp 2). The classified sound reflects the template nearest to the test trial in n-dimensional Euclidean space (*n* = PSTH bins). A confusion matrix reflecting predicted/actual outcomes for all trials is used to calculate accuracy, mutual information (Ml; bits) and encoding efficiency (bits/spk). **B.** Group data: sound classification accuracy between rules by unit type and depth, **a**. Accuracy in the context of V_R_. Left: Accuracy for individual units by rule. Right: Accuracy fold change between conditions (A-rule/V-rule); values <1 indicate lower A-rule accuracy (box plots as in Fig. 4). **b**. Accuracy in the context of V_U_ (conventions as in a.). **C.** Group data: encoding efficiency (bits/spike) between rules by unit type and depth, showing significant increases in deep BS units (conventions as in B). **a**. Encoding efficiency in the context of V_R_. **b**. Encoding efficiency in the context of V_U_.

We found that task rule could be decoded at greater than chance levels from responses to all four AV stimuli, and at all depth and NS/BS groups, showing that attentional state modulates decision stimulus PSTH responses throughout AC (Fig. S1; Table S5). These comparisons suggest response modulation by task rule, but do not address how information processing changes *across* the rules. To test this, we next used the decoder to compare accuracy in discriminating between responses to A_R_* (rewarded in A-rule) and responses to A_U_* (unrewarded in A-rule) bimodal stimuli across A-rule and V-rule conditions. This mimics the tone cloud discrimination required by the mice during the A-rule. In both rules, classification accuracy for the auditory decision stimuli (A_R_, A_U_) was higher than chance for all depth and BS/NS groups, regardless of whether the auditory stimuli were paired with V_R_ (**Fig. 6.D.a**; all p≤2.1e-05, all |z|≥4.2, one-way WSR vs chance [50%]; see **Table S6.A** for stats) or V_U_ (**Fig. 6.D.b**; all p<2.3e-06, all z > 4.7, one-way WSR vs chance [50%]; see **Table S6.B** for stats). Sound classification accuracy (A_R_*, A_U_*) did not significantly differ *across* the A-rule and V-rule (**Fig. 6.D.a**, A_R_V_R_ *vs.* A_U_V_R_ *comparison across rules:* all p≥0.38, all z≤0.88, see **Table S7.A** for full stats; **Fig. 6.D.b**; A_R_V_U_ *vs.* A_U_V_U_: all p≥0.14, all |z|≤1.46, see **Table S7.B** for full stats; paired WSR on decoder accuracy in A-rule vs. V-rule). Despite a reduction in activity levels during auditory attention, there was no loss in decoder accuracy, suggesting a possible change in encoding efficiency.

Through analysis of all decoder runs, we found that classifier accuracy and information were indeed correlated with FR (accuracy: r(3001) = 0.49, p = 2.3e-180; MI: r(3001) = 0.41, p = 1.5e-123; Pearson’s correlation, all A_R_* vs A_U_* decoder runs). Thus, normalizing information by mean joint per-trial spike rate for the two responses in each decode (bits/spike) provides insight into the efficiency with which spikes are used to represent stimuli. We found that this encoding efficiency measure increased by ~20% during the A-rule for deep-layer BS units, which was consistent between rewarded (A_R_V_R_ vs A_U_V_R_) and unrewarded (A_R_V_U_ vs A_U_V_U_) visual stimulus contexts (**Fig. 6.E.a**, A_R_V_R_ *vs.* A_U_V_R_ *comparison across rules:* Deep BS: p = 9.2e-05, Z = −3.91, paired WSR; med. FC: 1.26 [fold change: A-rule/V-rule]; V-rule: 0.11 ± 0.10, A-rule: 0.14 ± 0.15, mean bits/spk ± SDs, n = 147; all other groups p≥0.15, all |Z|≤1.44; see **Table S8.A** for full stats; **Fig. 6.E.b**, A_R_V_U_ *vs.* A_U_V_U_ Deep BS: p = 0.0086, Z = −2.63, med. FC: 1.19, V-rule: 0.13 ± 0.22, A-rule: 0.14 ± 0.13, n = 168; all other groups p≥0.22, all |Z|≤1.23; see **Table S8.B** for full stats). No other unit subpopulations showed significant changes. These outcomes provide important context for interpreting net spiking reductions during the A-rule by suggesting such changes serve to increase efficiency of sound encoding in deep layer BS units.

### Information encoding efficiency changes are driven by suppressed units

The finding that A-rule encoding efficiency increased and average FRs decreased in the deep layer BS unit group led us to further explore the relationship between activity level and information changes. Specifically, we sought to explore whether group-level information efficiency changes are driven by SUs with suppressed responses, and how the minority of units with increased A-rule FRs perform in the decoder. We therefore examined classifier accuracy and encoding efficiency for target and distractor (A_R_* vs A_U_*) decoding separately for deep-layer BS units with increased (**Fig. S2.B-C**) and decreased (**Fig. S2.B-D**) FRs in the A-rule. We found that units with increased FRs (36%; n = 65) exhibited a significant increase in A-rule decoding accuracy and MI (**Fig. S2.C.b;** p = 0.0042, Z = −2.87, med. FC: 1.05, V-rule % correct: 73.14 ± 17.00, A-rule: 77.01 ± 14.25, n = 66; paired WSR; fold change: A-rule/V-rule; mean ± SDs), but no significant change in encoding efficiency (**Fig. S2.C.c**; p = 0.31, Z = 1.02, V-rule bits/spk: 0.17 ± 0.20, A-rule: 0.14 ± 0.12). By contrast, units with suppressed FRs (64%; n = 114) showed no significant change in decoding accuracy (**Fig. S2.D.b**; p = 0.13, Z = 1.53, V-rule % correct: 71.64 ± 15.71, A-rule: 70.13 ± 15.17), but a 45% increase in encoding efficiency (**Fig. S2.D.c**; p = 3.7e-08, Z = −5.50, med. FC: 1.45, V-rule bits/spk: 0.09 ± 0.09, A-rule: 0.14 ± 0.13; paired WSR). These results suggest that the minority of units that increase FR in the A-rule perform marginally better at decoding the auditory stimulus, and that the units that decrease FR drive the shift in encoding efficiency.

### Receptive fields mapped during the inter-trial interval also show increased stimulus encoding efficiency

The analyses above revealed that auditory-selective attention reduced SU activity levels in AC, and that reduced spiking evoked by decision sounds reflected increased encoding efficiency. We next used another information theoretic approach to determine whether reduced spiking responses to behaviorally irrelevant sounds similarly reflected changes in information. In contrast to the discrete, repeated tone clouds used for decision stimuli, sounds presented during ITIs were continuous, non-repeating RDS segments. Reverse correlation was used to generate two spectrotemporal receptive fields (STRFs) for each SU: one from RDS stimuli presented during the A-rule and one from the V-rule (**Fig 7.A**). STRFs reflect the average stimulus energy preceding a spike and reveal features in both temporal and spectral domains that increase (red) or decrease (blue) spiking relative to the mean driven rate. Only units with a significant STRF in at least one attentional condition were included in further analyses, determined by a response reliability metric comparing the observed STRF to a null STRF obtained using time-reversed stimuli (**Fig. 7.B-C**). Significant STRFs were obtained in a fraction of all recorded units (143 BS, 58 NS), likely due to limited stimulus-response sampling; between each trial, 3.0 ± 2.1 s (median ± SD) of RDS stimulus was presented, with each duration determined by locomotion-based advancement to the next trial.

**Fig. 7.**
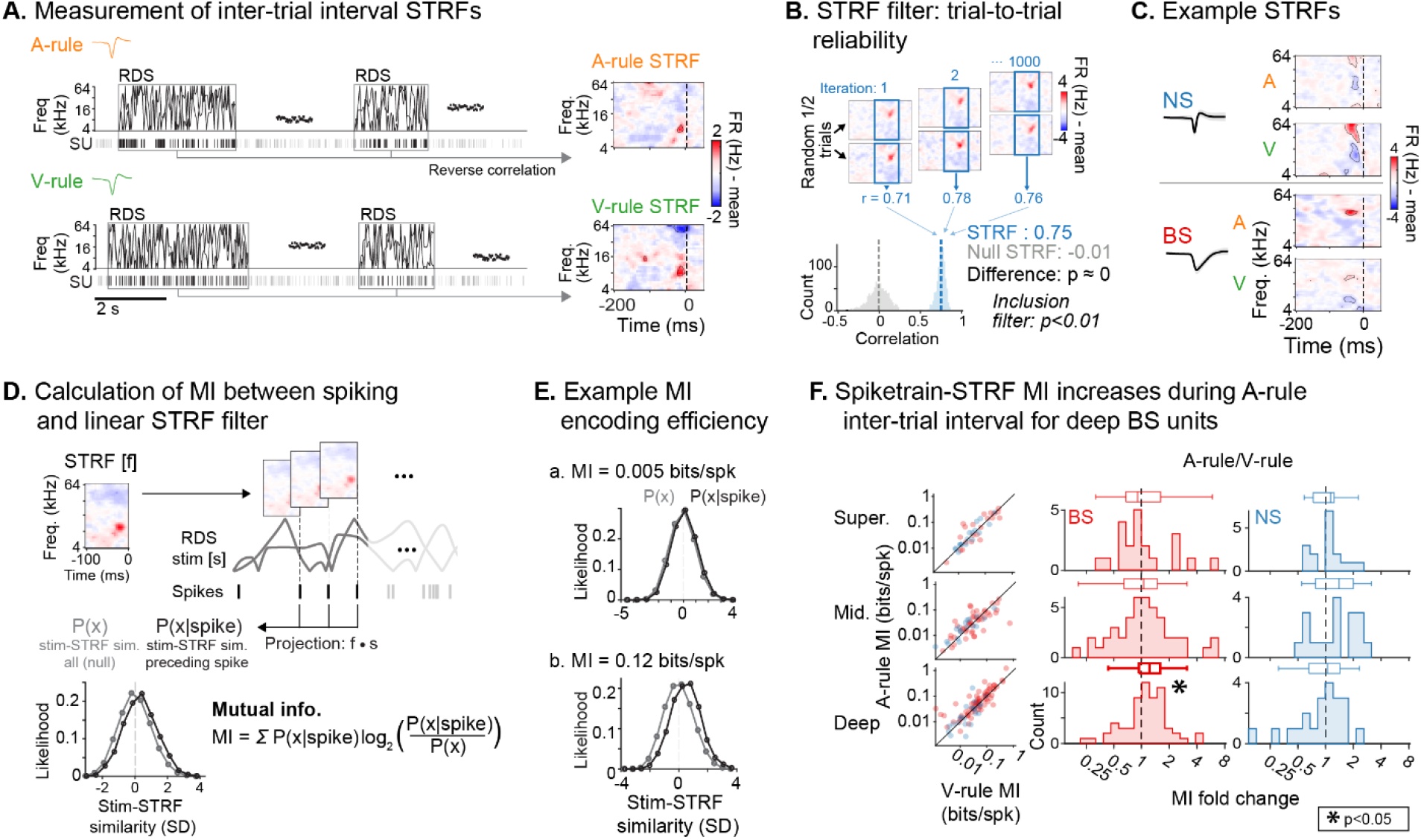
Auditory attention modulates inter-trial stimulus encoding efficiency. **A.** STRFs calculated separately for each condition by reverse correlation using spikes driven by the intertrial RDS stimulus. **B.** STRF significance determined by response reliability. Top: Correlations between STRFs calculated from subsampled data halves. Analysis window indicated in blue. Bottom: Correlation distributions form n = 1000 subsampled STRFs and Null STRFs (calculated using circularly-shifted spike trains). **C.** Example SU STRFs by rule. **D.** Encoding efficiency estimation. The STRF is convolved with the stimulus to define distributions of relative STRF-stimulus similarity values for all binned stimulus time points (*P(x)*) and those preceding a spike (*P(x|spike)*). Encoding efficiency reflects divergence between these distributions, which increases when spiking preferentially occurs during periods of higher stimulus-STRF similarity. **E.** Spiketrain-STRF encoding efficiency: low **(a)** and high **(b)** bits/spk examples. **F.** Group data: encoding efficiency (bits/spike) between rules by unit type and depth, showing significant increases in deep BS units (conventions as in Fig. 6). Left: Encoding efficiency for individual units by rule. Right: Encoding efficiency fold change between conditions (A-rule/V-rule); values >1 indicate increased A-rule encoding efficiency.

To calculate MI between the STRF and the spiketrain for each SU, we first defined probability distributions of STRF-stimulus projection values for all stimulus time points (*P*(*x*)) and for those time points preceding a spike (*P*(*x|spike*); Fig. 7.D). Intuitively, these projection values reflect the similarity between a windowed stimulus segment at a given timepoint and the STRF. The divergence of the two projection distributions is captured in a spike-rate normalized MI measure (bits/spk; encoding efficiency), which describes the reliability with which spikes are determined by stimulus features of the STRF. No differences in encoding efficiency between conditions were observed in the superficial or middle BS/NS groups, or the deep NS group. Instead, consistent with our earlier findings for decision stimuli, encoding efficiency showed a significant A-rule increase in the deep BS subpopulation (**Fig. 7.F**; Deep BS: p = 0.011, Z = −2.54, med. FC: 1.17, n = 73; paired WSR; fold change: A-rule/V-rule; mean bits/spk ± SDs; all other groups p≥0.31, all |z|≤1.02; see **Table S9** for full stats). This finding shows that auditory attention results in encoding of stimuli that is better described by a linear STRF filter and thus better tracks physical sound features. Furthermore, it suggests that increased encoding efficiency resulting from decreased spiking is a general effect of auditory attention in deep layer BS units regardless of the context-based behavioral relevance or learned valence of the sounds.

### Attention-related firing rate changes predict correct task performance

Analyses above revealed net suppression of pre-stimulus and evoked FRs during correct A-rule trials. If such modulation is behaviorally meaningful, we hypothesized that FRs preceding A-rule error trials may resemble sound-unattended V-rule trials. We addressed this possibility by comparing pre-stimulus FRs in error versus correct trials (300 ms prior to stimulus onset; **Fig. 8.B**). Because misses were uncommon (**Fig. 8.A**), we restricted our analysis to the comparison of FA and CR trials to allow for adequate sampling of each trial outcome. We included only behavior sessions with at least 10 FA and CR trials (A-rule and V-rule trials considered separately). When considering all BS units, we found no significant group-level difference between A-rule FA and CR trials at any cortical depth (**Fig. 8C; Table S10.A** for full stats; paired WSR: all p≥0.13, all |Z|≤1.50). However, BS units with A-rule suppression (the dominant group level change across rules) showed significantly higher pre-stimulus FRs prior to A-rule FA trials than CR trials across depths (**Fig. 8C**; Super. BS [n = 31]: mean FR difference between pre-stim FA and CR trials = 0.94 Hz, p = 0.0034, Z = −2.93; Mid. BS [n = 45]: mean difference = 0.49 Hz, p = 0.014, Z = −2.45; Deep BS [n = 139]: mean difference = 0.26 Hz, p = 0.0052, Z = −2.80; paired WSR; see **Table S10.A**). This outcome was specific to the A-rule: pre-stimulus FRs in A-rule-suppressed units did not differ between FA and CR trials in the V-rule (**Table S10.B**; paired WSR: all p≥0.22, all |z|≤1.06). A-rule suppressed NS units similarly showed elevated pre-stimulus FRs on FA trials during the A-rule, but only for the superficial subpopulation (**Fig. 8C**: Super. NS [n = 6]: mean FR difference between pre-stim FA and CR trials = 3.5 Hz, p = 0.043, Z = −2.02; all other p≥0.25, |z|≤1.16; see **Table. S10.A**). An important caveat is that unit inclusion criteria resulted in small NS unit samples, raising the possibility that this analysis was underpowered to detect significant error-predictive FR changes in middle and/or deep layer subpopulations. Together, these findings suggest that FR reductions typical of modality-selective attention directly relate to behavioral outcomes.

**Fig. 8.**
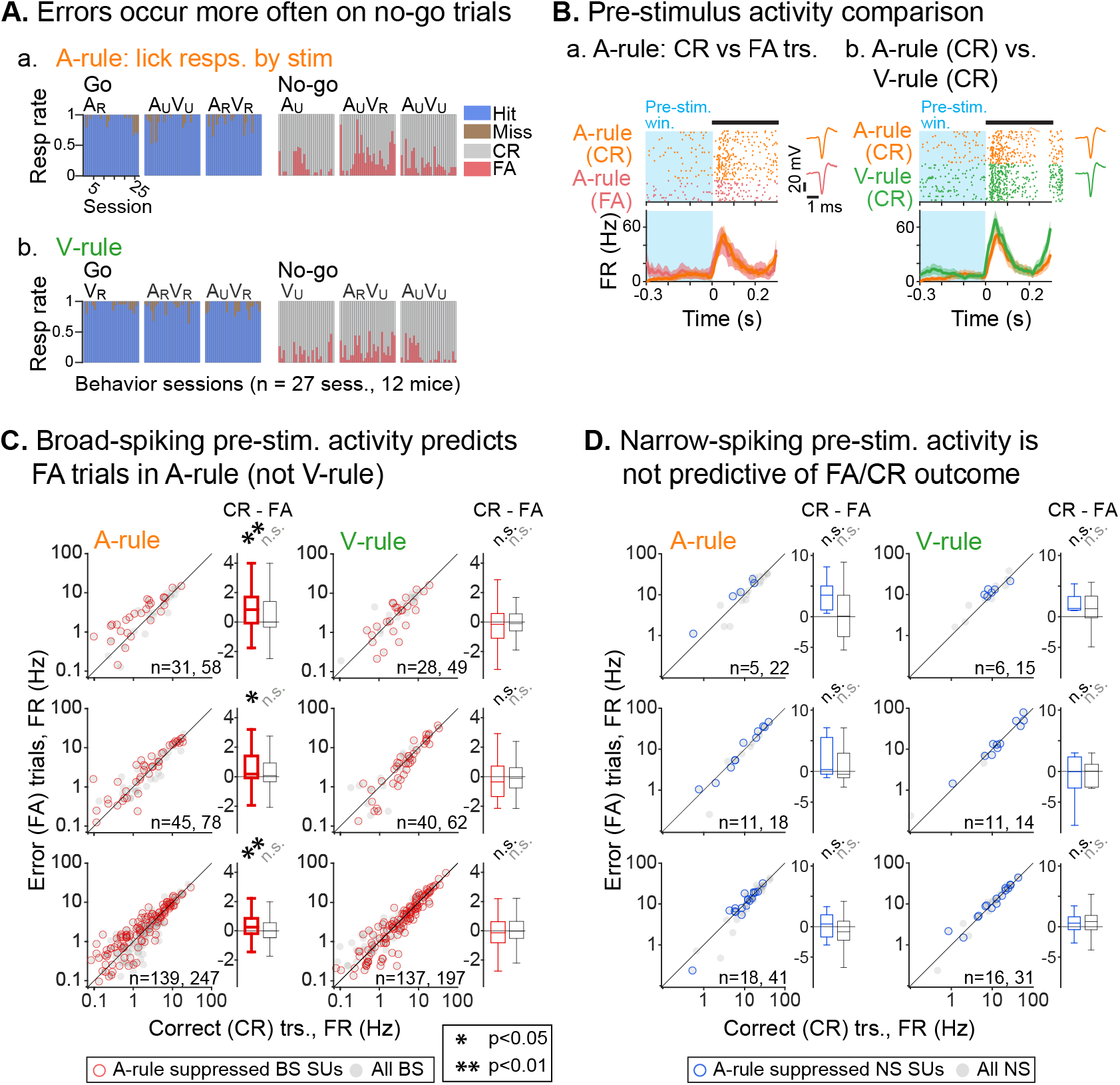
Attentionally suppressed broad-spiking units predict sound discrimination during auditory attention. **A.** Summary of behavioral outcomes for each session by stimulus (left/right) and rule (top/bottom). Errors are predominantly false alarms. **B.** Example unit with accuracy- and rule-dependent FR responses. Pre-stimulus FR analysis window shown in blue (−0.3 to 0 s). **a.** FRs elevated for FA compared to CR trials during A-rule. **b.** FRs on CR trials elevated during V-rule compared to A-rule. **C.** Group data: Pre-stimulus FRs in BS units between CR and FA trials by depth and rule (units with ≥10 FA and CR trials included). A-rule suppressed BS units show increased FRs on FA trials during A-rule but not V-rule. Box plots indicate FR difference values (CR - FA trials, Hz). **D.** Group data for NS units (conventions as in C). A-rule suppressed Superficial NS units show increased FRs on FA trials during A-rule.

## Discussion

In the present study, we recorded SU activity across AC layers in mice performing an audiovisual rule-switching task. We show that attention to sound results in a shift in AC stimulus representation marked by decreased activity and increased encoding efficiency, and that deep AC is a locus of such modulation. This activity reduction is behaviorally meaningful, as error trials are predicted by higher firing rates in the set of units that is suppressed in the A-rule. Many previous studies examining behavioral modulation of AC activity have relied on comparisons between task-engaged and passive exposure conditions (Bagur et al., 2018; Carcea et al., 2017; Kuchibhotla et al., 2017; Otazu et al., 2009), which may differ in terms of arousal, attention, reward expectation, and motor activity (Saderi et al., 2021). By holding arousal, motor activity, and physical stimulus features constant between conditions, we were able to isolate attentional effects on AC activity. The rule-switching task was difficult for the mice, considering over one third of sessions did not meet performance criteria (d’ >1.5 in both conditions). Indeed, until recently, studies of modality-selective attention were primarily conducted using human and nonhuman primate subjects. The ability to study such processes in a genetic model such as the mouse may open the door for further work on attentional modulation in cell-type specific microcircuits.

Classic work on feature selective attention suggests multiple mechanisms for highlighting behaviorally relevant signals. Attended features often evoke increased FRs, thereby increasing the reliability of sensory cortical readout (Desimone and Duncan, 1995; Moran and Desimone, 1985; Reynolds and Chelazzi, 2004). Noise reduction mechanisms may operate in tandem with such target enhancement, including distractor suppression (Schwartz and David, 2018), reduced spontaneous activity (Buran et al., 2014), reduced FR variability (Mitchell et al., 2007), and decreased noise correlations in neuronal populations (Cohen and Maunsell, 2009; Downer et al., 2015). In the present study, net reductions were observed in both pre-stimulus and evoked FRs during auditory attention. These changes were approximately balanced, such that baseline-divided FRs evoked by task decision sounds were largely unaffected by attention state. However, by accounting for spike timing patterns in a single-trial classifier analysis (Hoglen et al., 2018; Krishna and Semple, 2000; Malone et al., 2007), we found that stimulus information was preserved despite the loss of spikes, indicating increased encoding efficiency. Notably, auditory attention similarly reduced FRs evoked by task-irrelevant sounds, and encoding efficiency was again increased in STRF analysis accounting for spike timing patterns. Together, these findings suggest auditory attention imposes a general filter on information processing in AC, wherein spikes unrelated to sound encoding are differentially suppressed. This interpretation is consistent with an influential theory suggesting attention serves to suppress internally generated spontaneous activity in favor of processing behaviorally relevant external stimuli (Harris and Thiele, 2011).

As in previous studies, attention-related modulation was not uniformly expressed across cortical depths and neuron types. Changes in both FR and encoding efficiency were most prominent in deep layer neurons. These findings extend several previous studies reporting larger effects of attention in infragranular LFP and multi-unit activity (O’Connell et al., 2014; Zempeltzi et al., 2020). These physiological outcomes are consistent with anatomical work suggesting top-down modulatory signals arrive primarily in the supragranular and infragranular layers (Felleman and Van Essen, 1991). As the main cortical output layer, information shifts in the infragranular population would differentially influence subcortical sites and other cortical regions (Salin and Bullier, 1995). Previous studies also reported larger effects of task engagement in inhibitory interneurons. Although we observed larger fractions of NS units with significant modulation effects, group-level modulation was comparable between NS and BS units. Moreover, group-level increases in encoding efficiency were restricted to deep layer BS units.

In summary, we demonstrate a novel connection between attention-induced shifts in activity levels and stimulus encoding in early sensory cortex, which are directly related to behavioral outcomes. Previous research suggests such outcomes reflect top-down control by executive networks comprising frontal, parietal, thalamic, and striatal areas (Cools et al., 2004, 2006; Crone et al., 2006; Licata et al., 2017; Rikhye et al., 2018; Rougier et al., 2005; Toth and Assad, 2002; Wimmer et al., 2015). These networks may act as a context-dependent switch, routing attentional modulatory feedback to the appropriate sensory systems. In the present study, we provide evidence that such modulation specifically suppresses stimulus-irrelevant spiking, thus enhancing encoding efficiency in deep AC neurons.

## Methods

### Animals

All experiments were approved by the Institutional Animal Care and Use Committee at the University of California, San Francisco. Twenty-seven C57BL/6 background male mice were surgically implanted with a headpost and began behavioral training, of which 12 completed the training and successfully performed the task during physiology recording sessions. All mice began the experiment between ages P56 and P84. Mice used in this report expressed optogenetic effectors in a subset of interneurons, which we intended to use for optogenetic identification of cells (Lima et al. 2009; analysis not included here). These mice were generated by crossing an interneuron subpopulation-specific Cre driver line (PV-Cre JAX Stock Nr. 012358; Sst-Cre: JAX Stock Nr. 013044) with either the Ai32 strain (JAX Stock Nr. 012569), expressing Cre-dependent eYFP-tagged channelrhodopsin-2, or the Ai40 strain (JAX Stock Nr. 021188), expressing Cre-dependent eGFP-tagged archaerhodopsin-3. Of the 12 behavior mice included in this report, 7 were Ai32/Sst-Cre, 4 were Ai32/PV-Cre and one was Ai40/Sst-Cre. In some experiments, brief, low-level optogenetic pulses during the inter-trial interval of the task were used to identify opsin-expressing neurons (<0.3 mW light; 5 light pulses of 10 ms duration, every ~1.5 min); analysis of these results are outside of the scope of this report.

All mice were housed in groups of 2-5 for the duration of the behavioral training until the craniotomy. Post-craniotomy and during physiology recordings mice were housed singly (up to 6 days) to protect the surgical site. Mice were kept in a 12 hr/12 hr reversed dark/light cycle. All training occurred during the dark period, when mice showed increased activity and behavioral task performance (Roedel et al. 2006).

### Audiovisual rule-switching behavior task

Adult mice were trained on an audiovisual (AV) go/no-go rule-switching behavior task. In this task, mice were positioned on a floating spherical treadmill in front of a monitor and a speaker, and an optical computer mouse recorded treadmill movement. Mice licked to receive a reward depending on auditory, visual or AV stimulus presentation (“decision” stimuli, comprising both target and distractor), but the modality predictive of the reward changed partway through the behavioral session. Each session would start with a unimodal go/no-go block, in which a series of auditory (A_R_, A_U_; 17kHz or 8 kHz tone clouds [TC]) or visual (V_R_, V_U_; upward or rightward moving gratings) stimuli was presented. After stimulus presentation, mice signaled choice by either licking a spout in front of the mouth or withholding licking. Licking at the go unimodal stimulus would trigger a water reward, while licking at the no-go would trigger a short dark timeout. After a fixed number of unimodal trials, the stimuli would become AV, but the rule for which stimulus predicted reward would carry over from the unimodal block. All four stimulus combinations (A_R_V_R_, A_R_V_U_, A_U_V_R_, A_U_V_U_) would be presented in the AV block, such that two AV combinations would be go stimuli and two would be no-go. Then, after completing a fixed number of trials in the AV block, the task using the rule of the opposite modality would begin; a unimodal block with the other modality would start, followed by a second AV block using the rule from the preceding unimodal block. For any mouse, the stimuli predictive of the reward in each rule was kept constant across days and training sessions (e.g., a 17 kHz TC would always predict a reward in the A-rule, and a rightward grating would always predict a reward in the V-rule). In most cases, blocks proceed without interruption, although in some cases, the task is paused and resumed momentarily if the mouse fails to correctly transition between rules.

The task was self-paced using a virtual foraging approach, in which mouse locomotion (measured through treadmill rotation) would cause a track of randomly placed dots on the screen to move down. After a randomly varied virtual distance, a decision stimulus would be presented, at which point the mouse would lick or withhold licking to signal choice. For receptive field mapping during physiology experiments, a random double-sweep (RDS) stimulus was presented in between decision stimuli, during the inter-trial track portion.

### Behavior training and apparatus

Prior to any training, mice were surgically implanted with a stainless steel headplate for head fixation during the task, and later, for physiology recordings (methods described below). Three days post-implant, mice began a water restriction protocol based on previously published guidelines (Guo et al. 2014). Throughout the course of training, mice received a minimum water amount of 25 mL/kg/day, based on weight at time of surgical implant. After recovery from surgery, mice were given ~7 days to adjust to water restriction. Then, mice were head fixed and habituated to the floating treadmill for 15-30 min daily sessions with no stimulus presentation for 2-3 days. After mice appeared comfortable on the treadmill, a phased behavioral task training regimen began. Mice were trained once daily for ~6 days per week. On day 1, mice were introduced to an auditory-only (A-only) “stimulus training” version of the task in which A_R_ (“go”/“rewarded”) or A_U_ (“no-go”/“unrewarded”) stimuli were presented, and a reward would be automatically administered shortly after the onset of A_R_. Next, the mice were put on an operant version of the A-only task, which required licking any time after the onset of A_R_ to receive a reward and withholding of licking during A_U_ to avoid a dark timeout punishment. Mice achieved proficiency, defined as two or more consecutive days of sensitivity index d’ > 1.5 (see ‘Data analysis’ for calculation), on the A-only task after 9.5 ± 3.0 d after start of training (median ± SD, n = 12 successful mice). Then, a similar training structure was repeated for the visual task: V-only stimulus training with automatic rewards for V_R_, but not V_U_, followed by an operant version of the visual task requiring licks for rewards (median time to proficiency: 24.0 ± 4.5 d after start). After learning the tasks for each modality separately, mice were introduced to an auditory-AV (A-AV) version, in which the rule from the auditory stimulus carried over to the AV block. This was intermixed with training days on a visual-AV (V-AV) version of the task. Number of training days on A-AV or V-AV were decided based on prior performance, with extra training given as needed. Mice were considered proficient at this stage after performing with d’>1.5 on each rule (A-AV; V-AV) on two consecutive days (median time to proficiency: 43.5 ± 7.0 d after start). Finally, the full rule-switching task was introduced (Fig. 1.D), generally alternating between days of V-rule-first and the A-rule-first task sequences but allocating more training days to task orders as needed. Because physiology recordings were acute and strictly limited to 6 days after craniotomy, we set a greater threshold for expert-level performance on the full task before advancing to physiology: three consecutive days of d’>2.5 (median time to expertise: 90.5 ± 25.8 d). Care was taken to train each mouse at a roughly consistent time of day (no more than ~1-2 hrs day-to-day variation). During expert-level task performance, mice typically completed 260-300 trials in a daily session (30 A-only; 100 to 120 A-AV; 30 V-only; 100 to 120 V-AV).

The behavior training setup was controlled by two computers: a behavior monitoring and reward control PC (OptiPlex 7040 MT, Dell) and a dedicated stimulus presentation machine running Mac OS X (Mac Mini, Apple). Both machines ran custom software built with MATLAB, and inter-machine communication used the ZeroMQ protocol. Auditory and visual stimuli were generated and presented using the Psychophysics Toolbox Version 3 (Kleiner, Brainard, and Pelli 2007). Water rewards were administered using a programmable syringe pump (NE-500, New Era Pump Systems, Farmingdale, NY), positioned outside of the sound-attenuating recording chamber. Early in training, water reward volume was set at 0.01 mL per correct response, but over training the reward volume was gradually decreased to 0.006 mL to achieve greater trial counts before the mouse reached satiety. Licking events were recorded using a custom photobeam-based lickometer circuit based on plans provided by Evan Remington (Xiaoqin Wang Lab, Johns Hopkins University). Licks were registered when an IR photobeam positioned in front of the lick tube was broken, queried at a sample rate of 100 Hz by an Arduino Uno microcontroller (Arduino, LLC).

### *In vivo* awake recordings during behavior

Animals in this experiment underwent two surgeries: first, before training a surgery to implant a custom steel headplate over the temporal skull using dental cement was conducted. The animal was anesthetized using isoflurane and a headplate was positioned over AC, ~2.5 mm posterior to bregma and under the squamosal ridge, to allow for physiology recordings after achieving task expertise. When mice completed the training regimen outlined above, a craniotomy surgery was performed. The animal was again anesthetized using isoflurane and an elliptical opening (0.75 mm wide x 1.5 mm long) was made in the skull over AC using a dental drill. This opening was promptly covered with silicone elastomer (Kwik-Cast, World Precision Instruments), and the animal was allowed to recover overnight. The following day, the animal was affixed by its headplate over the treadmill inside of a sound-attenuating recording chamber, the silicone plug over the craniotomy was removed and the craniotomy was flushed with saline. A silver-chloride ground wire was placed into the craniotomy well at a safe distance from the exposed brain. A 64-channel linear probe (20 μm site spacing; Cambridge Neurotech, Cambridge, UK) was slowly inserted in the brain using a motorized microdrive (FHC, Bowdoin, ME) at an approximate rate of ~1 μm / s (Fiáth et al. 2019). After reaching the desired depth, the brain was allowed to settle for 10 min, after which the water spout, lickometer, visual stimulus delivery monitor and speaker were positioned in front of the mouse, and the behavior session commenced. Behavior sessions were sometimes stopped early and restarted due to poor performance. In approximately half of behavior-physiology sessions (15 of 27 successful recordings), the task was stopped due to low performance after the rule transition and restarted at the beginning (unimodal block) of the second rule. To control for possible effects of task order, attempts were made to counterbalance recordings from A-rule first (18) and V-rule first (9) behavior sessions.

After completion of the behavior task, the water spout and lickometer were removed, and a series of auditory and/or visual passive experiments were conducted in order to characterize the response properties of the recording site. All stimuli were presented with the auditory and visual stimulation apparatus described above. Following completion of these experiments, the probe was slowly removed, and the brain was covered with a thin layer of freshly mixed 2.5% agarose in saline, followed by a layer of silicone elastomer. The animal was returned to its home cage, and the following day the physiological recording process was repeated. Recordings were made for up to 6 days after the craniotomy. The neural signal acquisition system consisted of an Intan RHD2000 recording board and an RHD2164 amplifier (Intan Technologies), sampling at 30 kHz.

### Auditory and visual stimuli

In-task auditory decision stimuli were 1 s TCs, consisting of 50 ms tone pips overlapping by 25 ms, with frequencies in a 1 octave band around either 17 kHz or 8 kHz. TCs were frozen for the duration of the task, so that each mouse always heard the same pip sequences, allowing for direct comparisons of sound-evoked neural responses across rules without concern that stimulus peculiarities may be driving observed differences. Visual decision stimuli consisted of a circular moving grating stimulus (33° diameter subtended visual space), which appeared at the center of the screen for 1 s (coincident with TC stimulus during bimodal presentation). Gratings moved either upward or rightward with a 4 Hz temporal frequency, 0.09 cycles/degree spatial frequency at 50% contrast. In between decision stimulus presentations, a random double-sweep (RDS) stimulus was presented for receptive field mapping (Gourévitch et al. 2015). The RDS comprised two uncorrelated random sweeps that vary continuously and smoothly between 4 and 64 kHz, with a maximum sweep modulation frequency of 20 Hz.

After the behavior task, passive auditory search stimuli (pure tones, click trains) were presented to characterize response properties of the electrode channel. Click trains consisted of broadband 5 ms white noise pulses, presented at 20 Hz for 500 ms duration. Pure tone stimuli consisted of 100 ms tones of varied frequencies (4 – 64 kHz, 0.2 octave spacing) and sound attenuation levels (30 – 60 dB in 5 dB linear steps), with an interstimulus interval of 500 ms.

Auditory stimuli were presented from a free-field electrostatic speaker (ES1, Tucker-Davis Technologies) driven by an external soundcard (Quad-Capture or Octa-Capture, Roland) sampling at 192 kHz. Sound levels were calibrated using a Brüel & Kjær model 2209 meter and a model 4939 microphone. Visual stimuli were presented on a 19-inch LCD monitor with a 60Hz refresh rate (Asus VW199), positioned 25 cm in front of the mouse and centered horizontally and vertically on the eyes of the mouse. Monitor luminance was calibrated to 25 cd/m^2^ for a gray screen, measured at approximate eye level for the mouse.

### Data analysis

#### Behavioral performance

Task performance was evaluated by calculation of the *d’* sensitivity index:

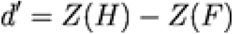

where *H* is hit rate and *F* is false alarm rate, and *Z* is the inverse normal transform. Because this transform is undefined for values of 0 or 1 and hit rates of 1 commonly occurred in this study, we employed the log-linear transformation, a standard method for correction of extreme proportions, for all calculations of *d’* (Hautus 1995). In this correction, a value of 0.5 is added to all elements of the 2 x 2 contingency table that defines performance such that:

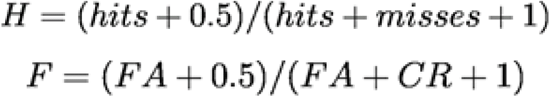

where *FA* is the false alarm count and *CR* is the correct reject count. To ensure that mice properly transitioned between task rules, *d’* values were calculated separately for responses in the A-rule and the V-rule. Behavioral sessions during physiological recording with *d’* < 1.5 in either rule were excluded from analyses (n = 28 successful sessions, n = 12 mice; 1 session excluded due to recording artifact, see below).

#### Spike sorting and unit stability evaluation

Spikes were assigned to unit clusters using KiloSort2 (KS2; Pachitariu et al. 2016). Clusters were first evaluated for isolation quality through the automated KS2 unit classification algorithm and then with a custom MATLAB interface. In this second step, clusters with non-neuronal waveforms or 2 ms refractory period violations >0.5% were removed from analysis (Laboy-Juárez et al., 2019; Sukiban et al., 2019). To evaluate stability, activity for each unit was plotted for the recording duration as a raster and binned spike counts (2 min bins) and manually examined for periods with a substantial drop off in FR (periods flagged for instability: 88 ± 10 % [mean ± SD] decrease in FR from median activity level). Flagged unstable periods were marked and removed from analysis (117 / 873 SUs with flagged durations >10% of recording time). Of the 28 physiology recording sessions with successful behavior, one was excluded due to a high degree of electrical noise contamination.

#### Classification of units by depth and waveform shape

Probes with electrode spans of 1260 μm were used, allowing for channels below and above AC. During recording, the probe was lowered to a point where several channels showed a prominent drop in field potential amplitude and spiking activity, indicating penetration into the white matter (Land et al. 2013). After behavior sessions, a set of auditory and visual stimulation protocols was used to map response properties of each electrode site, and multi-unit activity (MUA) responses were analyzed. Here, we define MUA as threshold crossings of 4.5 SD above a moving window threshold applied to each channel. Analysis of MUA was restricted to site characterization and is not included in the main results. We analyzed each tone or click PSTH for reliable responses, which we defined as trial-to-trial similarity of p<0.01 (Escabí et al., 2014). We designated the deepest channel with a reliable MUA sound response of any magnitude as the deep cortex-white matter border. Limited somatic spiking in the top layer of cortex prevented the use of MUA as a reliable marker for the superficial cortex-pia border (Senzai et al., 2019), so we instead relied on a LFP-based measure. To define the top border of cortex, the maximum spontaneous LFP (1-300 Hz) amplitude of a 10 s snippet from each channel was plotted, and the channel at which LFP amplitude dropped off to the approximate probe-wise noise floor (i.e., minimum LFP amplitude) was considered the top channel in cortex (**Fig. 3.B.c**). These measures were confirmed histologically through Di-I probe marking experiments with a separate group of untrained mice; histology methods described below and elsewhere (Morrill and Hasenstaub, 2018). Marking the top and bottom cortical borders generated a span of channels putatively within AC. This span was used to divide channels into superficial, middle, and deep groups, based on measurements of the fraction of cortex attributed to supragranular (layers 1-3), granular (layer 4) and infragranular (layers 5-6) in the mouse AC (Allen Institute Mouse Brain Atlas; https://mouse.brain-map.org/). SUs were assigned the fractional depth of the channel on which the largest magnitude waveform was recorded.

Clusters were also classified into broad-spiking (BS; putatively excitatory) and narrow-spiking (NS; putatively fast-spiking inhibitory) units on the basis of the bimodal distribution of waveform peak-trough durations (**Fig. 3.D;** NS/BS transition boundary = 0.6 ms). From sessions with successful behavior, we recorded 873 SUs from all cortical depths, comprising 17.2% (150) NS units and 82.8% (723) BS units.

#### Firing rate analysis and trial filters

To compare FR responses to stimuli across task rules and to the receptive field mapping stimulus, we measured FR in the first 300 ms post-stimulus onset. Only units with nonzero FRs in both rules were included. To ensure that measurements were capturing periods of task engagement, all trials with incorrect responses (misses and FAs) were excluded from all decision-stimulus analyses, with the exception of those shown in **Fig. 8**. We also excluded trials with recorded licks earlier than the 300 ms post-stimulus onset, or in the 500 ms pre-stimulus onset. Given these filters, analyses were restricted to units present in the recording during at least 10 trials (correct behavioral choice and without “early licks”) for each stimulus type.

#### PSTH-based Euclidean distance decoding

A peristimulus time histogram (PSTH)-based decoder was used to compare mutual information between A-rule and V-rule spiketrains and stimulus identity (**Fig. 6.A**; Foffani and Moxon, 2004; Hoglen et al., 2018; Malone et al., 2007). In this method, two or more responses are compared by generating template PSTHs by removing one test trial. This test trial response is also binned into a single-trial PSTH, and then classified as belonging to the nearest template in *n-*dimensional Euclidean space, where *n* is the number of PSTH bins. More formally, the nearest template is that which minimizes the Euclidean norm between test and template vectors (PSTHs). This process is then repeated for all trials comprising the template PSTHs. Decoding accuracy is the percentage of trial responses that are correctly assigned to the stimuli that elicited them. Mutual information (MI) is calculated from a confusion matrix of classifications as follows:

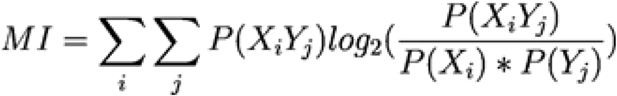

where *X* is the decoder prediction, *Y* is the actual, *P*(*X_i_ Y_j_*) represents the value of the (*i*, *j*) element of the confusion matrix, and *P*(*X_i_*) and *P*(*Y_j_*) aresums on the marginals. This yields a value of mutual information in bits. To measure encoding efficiency (bits per spike), we normalized mutual information by the joint mean spikes per trial of the responses submitted to the decoder (Bigelow et al., 2019; Buracas et al., 1998; Zador, 1998).

For consistency with FR analyses, a time window of 0 to 300 ms, where stimulus onset is 0, was chosen for decoding analysis. A PSTH binwidth of 30 ms was chosen based on optimal binwidth calculations for mouse AC using the same decoding method (Hoglen et al., 2018). To filter out units with low responsiveness to any of the stimuli in a given decoding analysis, we required a minimum FR of 5 Hz during the 0-300 ms window in at least one of the stimulus conditions. As such, unit sets may differ between each decoding analysis due to units that were responsive to one set of stimuli but unresponsive to others.

#### Spectrotemporal receptive field (STRF) analysis

To test whether task rule modulates auditory receptive fields, we presented a random-double sweep stimulus (RDS, described in *Auditory and visual stimuli*) in between trials for durations of ~1 to 15 s, depending on rate of task progression. Different randomly generated RDS segments were presented in each inter-trial interval, and STRFs were generated separately for each rule. Because total RDS duration varied between the A-rule and the V-rule in a single session, we equated presentation time across rules by truncating the segments of the rule with greater RDS time (presentation time in each rule: 7.58 ± 2.50 min [mean ± SD]; n = 27 sessions). This ensured that different stimulus presentation times did not bias STRF estimation. The first 200 ms of RDS response was dropped from all STRF analyses to minimize bias from strong onset transients. SU activity during these short RDS segments was used to generate STRFs for each segment using standard reverse-correlation techniques (Aertsen and Johannesma 1981). In brief, the spike-triggered average (STA) was calculated by adding all stimulus segments that preceded spikes using a window of 200 ms before and 50 ms after each spike. The choice of 200 ms prior to each spike reflects the upper limit of temporal integration times of auditory cortical neurons (Atencio and Schreiner 2013), and the 50 ms post-spike time was included to estimate acausal values, i.e., those that would be expected by chance given the stimulus and spike train statistics (Gourévitch et al. 2015). STRFs were transformed into units of firing rate (Hz) using standard methods discussed elsewhere (Rutkowski et al. 2002). Units with poorly-defined STRFs were filtered out using a trial-to-trial correlation metric (Escabí et al., 2014): STRF segments were randomly divided into two halves, re-averaged separately, and a correlation value was calculated for the two STRFs. This process was then repeated 1000 times, and the mean of correlations defined the reliability value for each STRF. We compared the mean observed STRF reliability to a null distribution of reliabilities, generated by repeating the procedure on null STRFs made from circularly-shuffled spike trains (preserving spike count and interspike interval but breaking the timing relationship between spikes and stimulus). A *p*-value was calculated as the fraction of the null STRF reliabilities greater than the mean observed STRF reliability, and STRFs with *p*<0.01 in either rule were included in subsequent analyses. Any STRFs from units with greater than 10% of recording duration marked as unstable using the aforementioned recording stability metrics were removed from analysis.

Mutual information between a spiketrain and an STRF was measured as the divergence of two distributions: one reflecting the similarity of the windowed stimulus segments (RDS) preceding a spike and the STRF, and the other reflecting the similarity of all possible windowed stimulus segments and the STRF, regardless of whether a spike occurred (**Fig. 7.D**; Atencio and Schreiner, 2012; Atencio et al., 2008; Escabí and Schreiner, 2002). Stimulus-STRF similarity was defined as the inner product of the STRF and the stimulus segment of equivalent dimensions, with higher values reflecting closer matches between the STRF and stimulus. The distribution *P*(*z|spike*) was generated from *z* = *s · STRF*, where *s* represents all RDS stimulus segments that preceded a spike. Then the distribution *P*(*z*) was made from similarity calculations of all possible windowed RDS segments and the STRF. The mean *μ* and the standard deviation (SD) *σ* of *P*(*z*) were calculated, and the distributions were transformed into units of SD: *x* = (*z – μ*)/*σ*, yielding distributions of *P*(*x|spike*) and *P*(*x*) expressed in units of SD.

Using the distributions described above, a spike count-normalized measure of mutual information between the calculated STRF and the spike train can be calculated as:

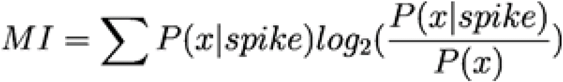

We used this value to compare how well STRFs from A-rule and V-rule ITIs predict a spiketrain, and thus whether activity in each attentional condition is well described by this canonical filter model.

### Statistics

All statistical calculations were performed in MATLAB r2019a and its Statistics and Machine Learning Toolbox, V_R_1.5. For comparisons of SU responses across task rules at the group level, a paired Wilcoxon signed-rank (WSR) test was used. To determine if individual SUs were significantly modulated by rule, an unpaired Student’s t-test on FR was used with a threshold of *p*<0.01. Descriptive statistics reported in text are mean ± standard deviation (SD), unless otherwise noted. Fractional change values between task rules are reported as the median of the A-rule/V-rule. All other statistical tests are described in Results. Sample sizes (*n*) are indicated for each comparison in Results or Supplemental tables.

### Histological verification of depth measurement

To test the accuracy of our depth estimation method based on physiological responses (Fig. 3), we presented the pure tone search stimuli described above to a separate set of untrained control mice while performing extracellular recordings (n = 11 recordings from 4 mice; Ai32/Sst-Cre). Before insertion, the probe was painted with the fluorescent lipophilic dye Di-I (DiCarlo et al., 1996; Morrill and Hasenstaub, 2018). The depth measurement procedure based on physiological signals was carried out as described above, and then probe tracks from each recording were visualized as described previously (Morrill and Hasenstaub, 2018). Briefly, after recordings, the animal was euthanized, and the brain was removed and placed into a solution of 4% PFA in PBS (0.1 m, pH 7.4) for 12 h, followed by 30% sucrose in PBS solution for several days. The brain was then frozen and sliced using a sliding microtome (SM2000R, Leica Biosystems) and slices were imaged with a fluorescence microscope (BZ-X810, Keyence). Di-I probe markings showing cortical depth were consistent with physiological activity-based depth measurements described above (Fig. 3.B-C).

## Acknowledgements

This work was supported by National Institutes of Health grant R01DC014101 to A.R.H., the National Science Foundation GRFP to R.J.M., the Klingenstein Foundation to A.R.H., Hearing Research Inc. to A.R.H., and the Coleman Memorial Fund to A.R.H.

## Supplemental materials

**Fig. S1.**
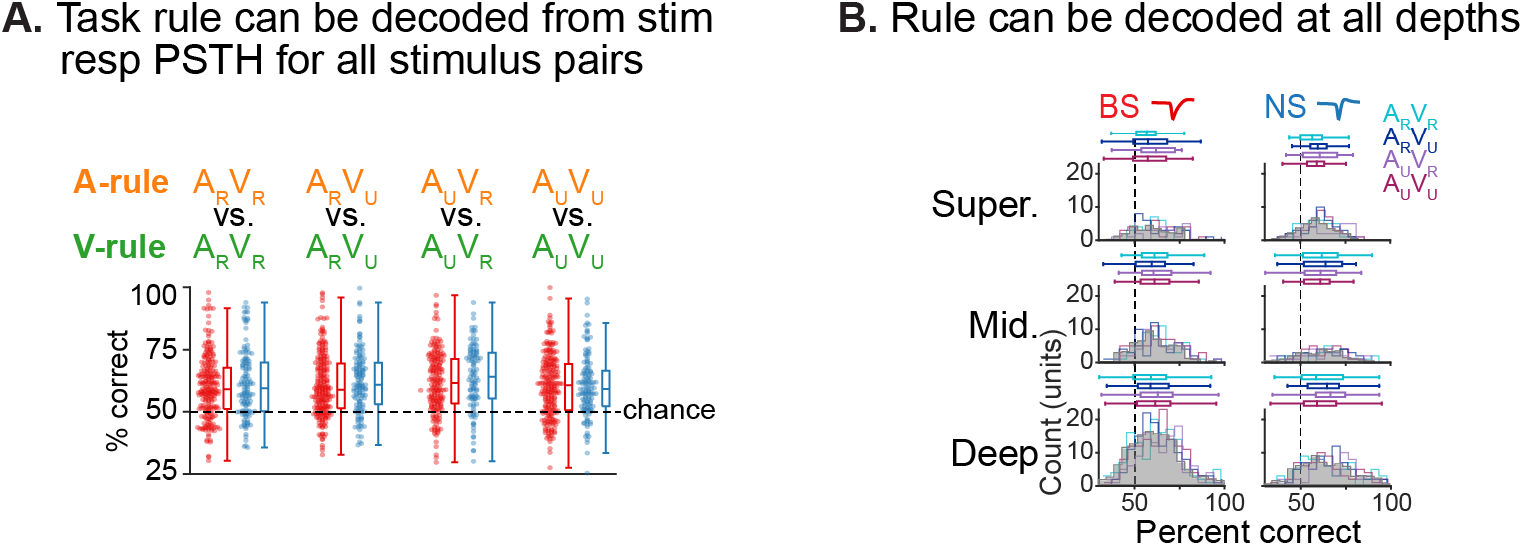
Decoding of task rule from PSTHs. **A.** PSTH-based Euclidean distance decoding of rule identity. Responses to each of the four AV stimuli in the A-rule vs responses to the same stimuli in the V-rule were decoded based on 0-0.3 s stimulus onset window; PSTHs were constructed with 30 ms bins. Each dot represents SU decoding accuracy (% correct); BS units in red, NS in blue. Chance decoding is indicated by the dashed line. A minimum 5 Hz FR response to at least one stimulus in the decode was required for inclusion. Box plots as before: central line indicates median, box edges indicate 25th and 75th percentiles, whiskers extend to data points not considered outliers. **B.** Decoding by depth group. Median decoding by stimulus indicated as colored horizontal box plots (as described above). Gray distributions represent decoding by depth, averaged across stimulus types. Left: broad-spiking units, right: narrow-spiking. See Table S5 for statistics.

**Fig. S2.**
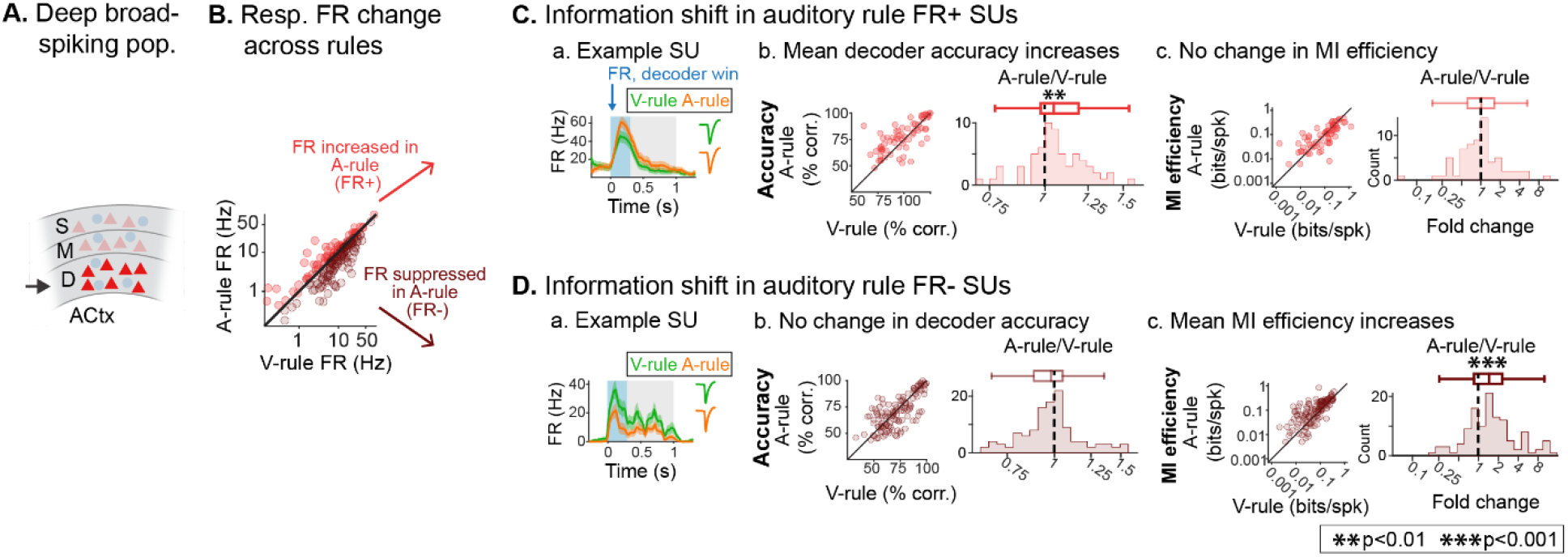
Deep AC A-rule information efficiency increases are driven by FR suppressed units. **A.** Analysis is restricted to unit group exhibiting increased information efficiency: deep broad-spiking units. **B.** Decoder-based analyses are split into subpopulations showing increased stimulus responses in the A-rule (FR+; n = 65, 36%) or decreased responses in the A-rule (FR-; n = 114, 64%). **C.** Restriction of analysis to FR+ units. **a.** Example FR+ unit. **b.** PSTH-based decoder accuracy for FR+ group shows a significant increase in A-rule. Conventions as in Fig. 4. Decoder accuracy shown is the mean of A_R_V_R_ vs. A_U_V_R_ and A_R_V_U_ vs. A_U_V_U_ decoding across rules. **c.** No change is observed for MI efficiency. **D.** Similar analyses for FR-units. **a.** Example FR-unit. **b.** Decoder accuracy does not change across rules despite loss of spikes in the FR-subpopulation. **c.** MI efficiency increases are driven by the FR-population, directly linking changes in information rate to changes in activity levels across attentional state.

**Table S1.A.** Analysis: A_R_* response FR across rules.

| *Group* | *n* | *A-rule*<br>*mean*<br>*FR (Hz)* | *±SD* | *V-rule*<br>*mean*<br>*FR (Hz)* | *±SD* | *WSR*<br>*Z-value* | *p-value* | *Median*<br>*fold*<br>*change*<br>*across*<br>*rules*<br>*(p<0.05*<br>*only)* |
| --- | --- | --- | --- | --- | --- | --- | --- | --- |
| Super. BS | 80 | 3.89 | 5.87 | 3.62 | 5.17 | -1.29 | 0.2 | -- |
| Super. NS | 35 | 14.75 | 14.63 | 15.78 | 17.28 | 1.03 | 0.3 | -- |
| Mid. BS | 153 | 6.44 | 13.48 | 6.42 | 9.4 | 2.55 | 0.011 | 0.87 |
| Mid. NS | 34 | 15.23 | 11.9 | 19.27 | 15.3 | 2.59 | 0.0096 | 0.8 |
| Deep BS | 391 | 5.17 | 7.8 | 6.02 | 8.74 | 4.7 | 2.7E-06 | 0.88 |
| Deep NS | 77 | 21.45 | 19.02 | 24.86 | 20.11 | 3.17 | 0.0015 | 0.87 |

**Table S1.B.** Analysis: A_U_* response FR across rules.

| *Group* | *n* | *A-rule*<br>*mean*<br>*FR (Hz)* | *±SD* | *V-rule*<br>*mean*<br>*FR (Hz)* | *±SD* | *WSR*<br>*Z-value* | *p-value* | *Median*<br>*fold*<br>*change*<br>*across*<br>*rules*<br>*(p<0.05*<br>*only)* |
| --- | --- | --- | --- | --- | --- | --- | --- | --- |
| Super. BS | 76 | 4.8 | 6.82 | 4.19 | 5.77 | -0.85 | 0.39 | -- |
| Super. NS | 35 | 15.75 | 12.36 | 17.29 | 13.66 | 1.17 | 0.24 | -- |
| Mid. BS | 148 | 5.7 | 10.73 | 5.83 | 7.57 | 2.42 | 0.015 | 0.81 |
| Mid. NS | 34 | 15.79 | 14.66 | 20.75 | 19.78 | 3.02 | 0.0025 | 0.76 |
| Deep BS | 374 | 4.87 | 7.98 | 6.02 | 10.06 | 5.56 | 2.7E-08 | 0.82 |
| Deep NS | 77 | 19.33 | 16.8 | 21.36 | 15.22 | 2.38 | 0.017 | 0.83 |

**Table S2.**
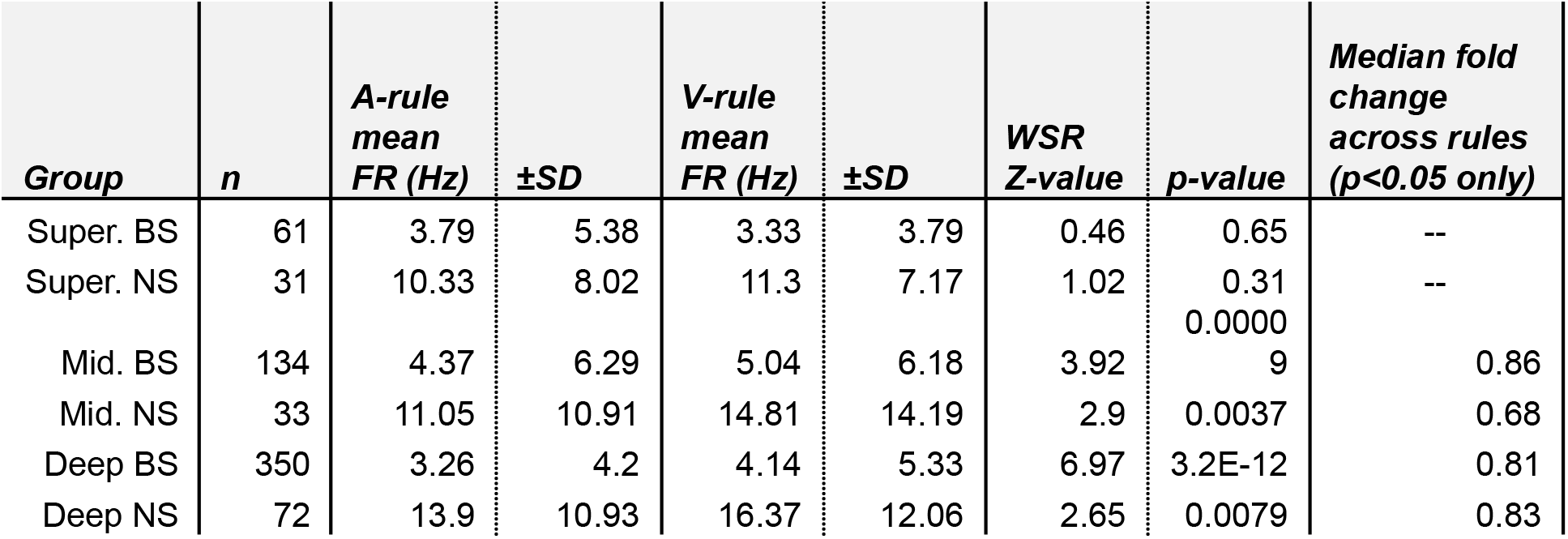
Analysis: RDS response FR across rules.

**Table S3.** Analysis: Pre-stimulus FR across rules.

| **Group** | ***n*** | ***A-rule***<br>***mean***<br>***FR (fold***<br>***change)*** | ***±SD*** | ***V-rule***<br>***mean***<br>***FR (fold***<br>***change)*** | ***±SD*** | ***WSR***<br>***Z-value*** | ***p-value*** | ***Median fold***<br>***change***<br>***across rules***<br>***(p<0.05***<br>***only)*** |
| --- | --- | --- | --- | --- | --- | --- | --- | --- |
| Super. BS | 82 | 3.27 | 5.05 | 2.62 | 3.38 | -1.5 | 0.13 | -- |
| Super. NS | 35 | 11.14 | 9.08 | 10.66 | 8.42 | -0.84 | 0.4 | -- |
| Mid. BS | 154 | 4.33 | 7.05 | 4.8 | 6.07 | 3.19 | 0.0014 | 0.88 |
| Mid. NS | 34 | 12.89 | 11.84 | 16.29 | 14.72 | 2.78 | 0.0055 | 0.76 |
| Deep BS | 403 | 3.4 | 4.49 | 4.2 | 5.54 | 5.47 | 4.4E-08 | 0.86 |
| Deep NS | 77 | 14.33 | 11.33 | 15.91 | 12.32 | 2.32 | 0.02 | 0.86 |

**Table S4.A.**
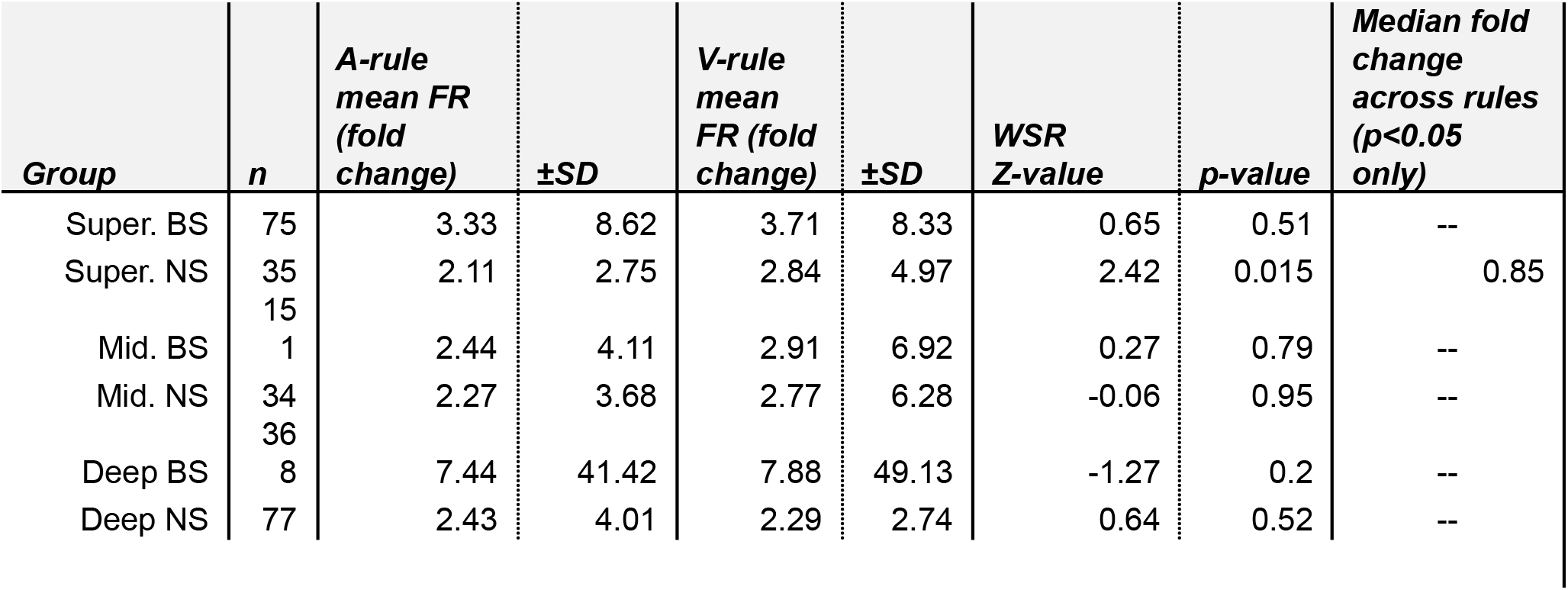
Analysis: A_R_* baseline-adjusted FR across rules.

**Table S4.B.**
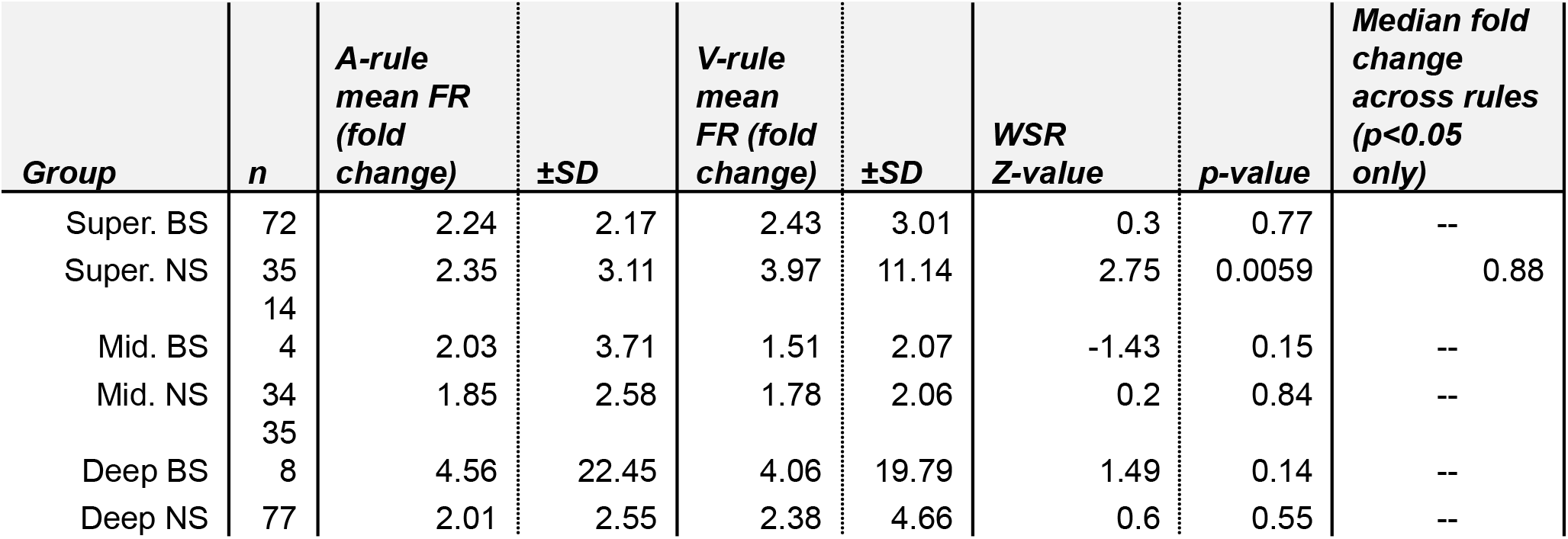
Analysis: A_U_* baseline-adjusted FR across rules.

**Table S5.A.** Analysis: decoder accuracy, A-rule vs V-rule by stimulus type Accuracy (%): mean ± SD.

| **Group** | **A<sub>R</sub>V<sub>R</sub>** | **A<sub>R</sub>V<sub>U</sub>** | **A<sub>U</sub>V<sub>R</sub>** | **A<sub>U</sub>V<sub>U</sub>** |
| --- | --- | --- | --- | --- |
| BS | 60 ± 12.9 | 60.7 ± 12.8 | 61.9 ± 12.5 | 60.6 ± 12.7 |
| NS | 61.2 ± 13.6 | 62.2 ± 12.5 | 63.7 ± 13.1 | 60.4 ± 12.4 |

One-way Wilcoxon signed rank (WSR) vs chance [50%]: all $p \leq 10^{-13}$ , all $|z| \geq 7.4$

**Table S5.B.** Analysis: decoder accuracy, A-rule vs V-rule by depth, BS/NS Accuracy (%): mean ± SD.

| **Group** | **Avg., all stim** |
| --- | --- |
| Super. BS | 58.5 ± 9.8 |
| Super. NS | 58.4 ± 7.1 |
| Mid. BS | 60.5 ± 9.2 |
| Mid. NS | 60.7 ± 10.2 |
| Deep BS | 61.0 ± 10.7 |
| Deep NS | 63.8 ± 11.2 |

One-way WSR vs chance [50%]: all $p \leq 10^{-4}$ , all $|z| \geq 4.3$

**Table S6.A.**
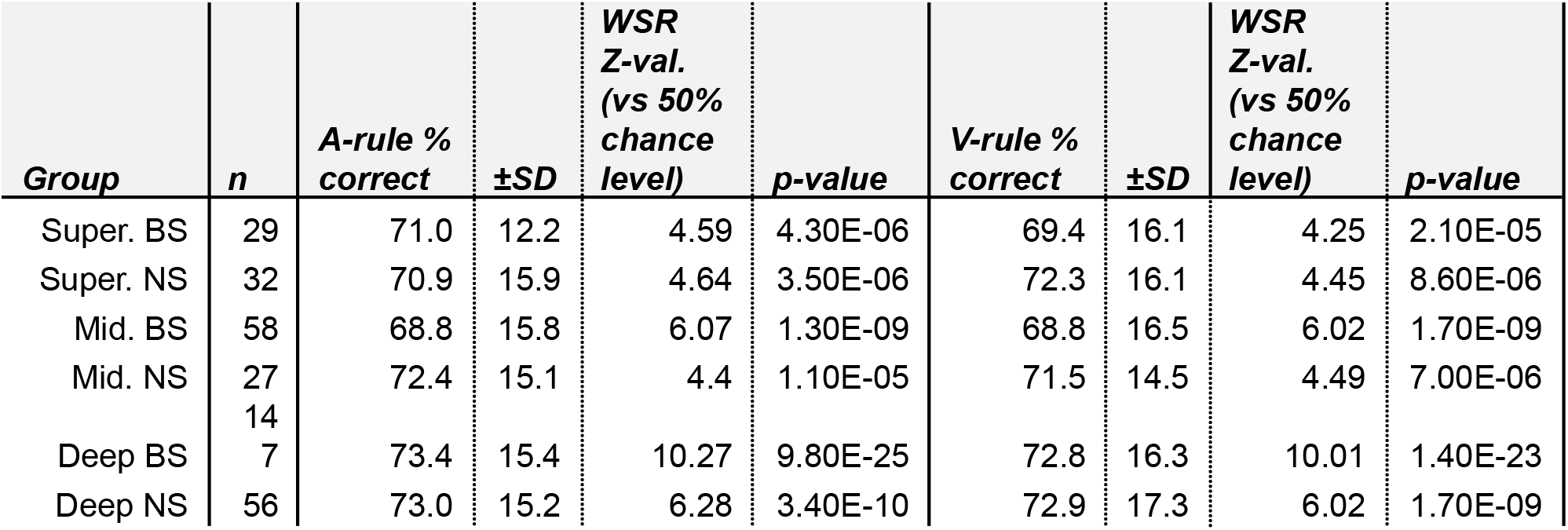
Analysis: decoder accuracy of A_R_V_R_ vs A_U_V_R_ across depth groups.

**Table S6.B.**
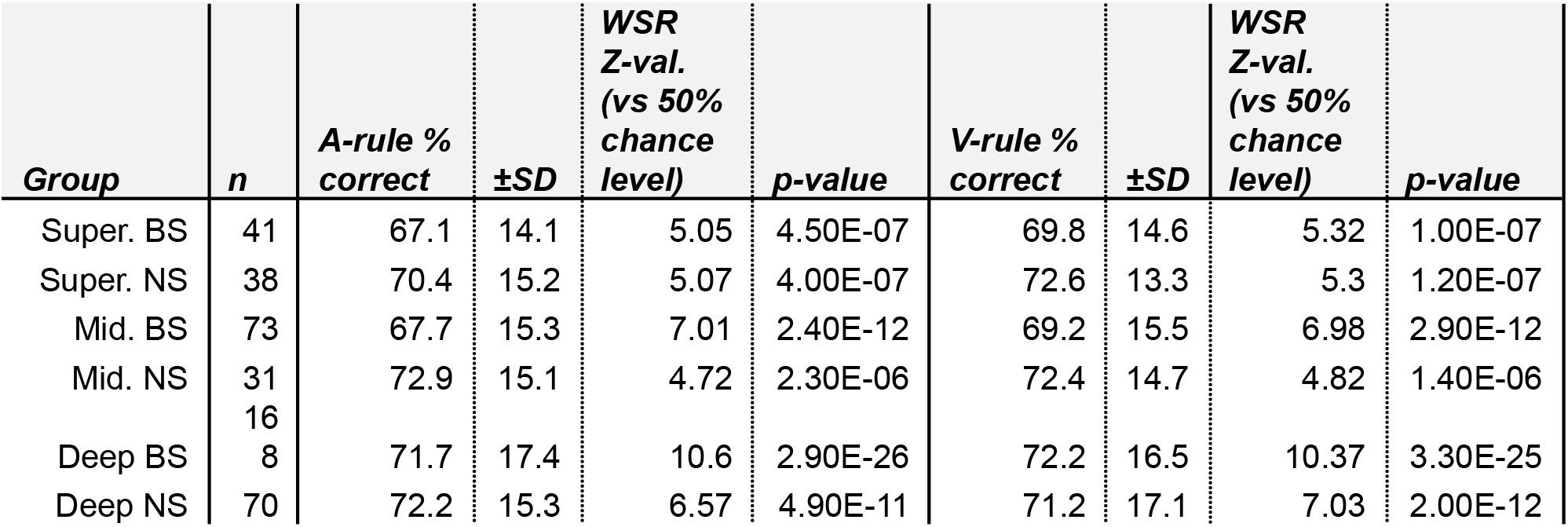
Analysis: decoder accuracy of A_R_V_U_ vs A_U_V_U_ across depth groups.

**Table S7.A.** Analysis: Wilcoxon signed-rank comparison of A_R_V_R_ vs A_U_V_R_ decoding across rules ** For data values, refer to table S6.A.

| **Group** | ***n*** | ***WSR Z-value***<br>***across rules*** | ***p-value*** |
| --- | --- | --- | --- |
| Super. BS | 29 | -0.88 | 0.38 |
| Super. NS | 32 | 0.71 | 0.48 |
| Mid. BS | 58 | 0.31 | 0.76 |
| Mid. NS | 27 | -0.53 | 0.60 |
| Deep BS | 147 | -0.49 | 0.62 |
| Deep NS | 56 | 0.51 | 0.61 |

**Table S7.B.** Analysis: Wilcoxon signed-rank comparison of A_R_V_U_ vs A_U_V_U_ decoding across rules ** For data values, refer to table S6.B.

| **Group** | ***n*** | ***WSR Z-value***<br>***across rules*** | ***p-value*** |
| --- | --- | --- | --- |
| Super. BS | 41 | 1.42 | 0.16 |
| Super. NS | 38 | 1.46 | 0.14 |
| Mid. BS | 73 | 1.01 | 0.31 |
| Mid. NS | 31 | -0.71 | 0.48 |
| Deep BS | 168 | -0.42 | 0.67 |
| Deep NS | 70 | 0.1 | 0.92 |

**Table S8.A.** Analysis: MI efficiency of A_R_V_R_ vs A_U_V_R_ across rules.

| *Group* | *n* | *A-rule*<br>*mean*<br>*bits/spk* | *±SD* | *V-rule*<br>*mean*<br>*bits/spk* | *±SD* | *WSR*<br>*Z-value*<br>*across*<br>*rules* | *p-val* | *Median*<br>*fold*<br>*change*<br>*across*<br>*rules (sig.*<br>*only)* |
| --- | --- | --- | --- | --- | --- | --- | --- | --- |
| Super. BS | 29 | 0.15 | 0.13 | 0.12 | 0.11 | -1.44 | 0.15 | -- |
| Super. NS | 32 | 0.09 | 0.09 | 0.08 | 0.07 | -0.82 | 0.41 | -- |
| Mid. BS | 58 | 0.12 | 0.13 | 0.12 | 0.13 | -0.27 | 0.79 | -- |
| Mid. NS | 27 | 0.11 | 0.24 | 0.07 | 0.08 | -0.41 | 0.68 | -- |
| Deep BS | 147 | 0.14 | 0.15 | 0.11 | 0.1 | -3.91 | **9.20E-05** | 1.26 |
| Deep NS | 56 | 0.06 | 0.06 | 0.05 | 0.05 | -0.56 | 0.57 | -- |

**Table S8.B.** Analysis: MI efficiency of A_R_V_U_ vs A_U_V_U_ across rules.

| *Group* | *n* | *A-rule*<br>*mean*<br>*bits/spk* | *±SD* | *V-rule*<br>*mean*<br>*bits/spk* | *±SD* | *WSR*<br>*Z-value*<br>*across*<br>*rules* | *p-val* | *Median*<br>*fold*<br>*change*<br>*across*<br>*rules (sig.*<br>*only)* |
| --- | --- | --- | --- | --- | --- | --- | --- | --- |
| Super. BS | 41 | 0.11 | 0.13 | 0.12 | 0.13 | 0.89 | 0.37 | -- |
| Super. NS | 38 | 0.08 | 0.09 | 0.06 | 0.06 | 0.17 | 0.87 | -- |
| Mid. BS | 73 | 0.1 | 0.09 | 0.09 | 0.09 | -0.31 | 0.76 | -- |
| Mid. NS | 31 | 0.07 | 0.07 | 0.06 | 0.08 | -0.78 | 0.43 | -- |
| Deep BS | 168 | 0.14 | 0.13 | 0.13 | 0.22 | -2.63 | **0.0086** | 1.19 |
| Deep NS | 70 | 0.06 | 0.06 | 0.05 | 0.05 | -1.23 | 0.22 | -- |

**Table S9.** Analysis: Comparison of STRF MI (bits/spk) across rules.

| ***Group*** | ***n*** | ***A-rule***<br>***mean***<br>***bits/spk*** | ***±SD*** | ***V-rule***<br>***mean***<br>***bits/spk*** | ***±SD*** | ***WSR Z-value***<br>***across rules*** | ***p-val*** | ***Median***<br>***fold***<br>***change***<br>***across***<br>***rules***<br>***(sig.***<br>***only)*** |
| --- | --- | --- | --- | --- | --- | --- | --- | --- |
| Super. BS | 22 | 0.1 | 0.1 | 0.1 | 0.11 | -0.23 | 0.82 | -- |
| Super. NS | 17 | 0.05 | 0.06 | 0.05 | 0.06 | -1.02 | 0.31 | -- |
| Mid. BS | 48 | 0.06 | 0.08 | 0.08 | 0.12 | 0.42 | 0.67 | -- |
| Mid. NS | 19 | 0.03 | 0.03 | 0.03 | 0.05 | -0.94 | 0.35 | -- |
| Deep BS | 73 | 0.12 | 0.19 | 0.1 | 0.18 | -2.54 | 0.011 | 1.17 |
| Deep NS | 22 | 0.03 | 0.05 | 0.03 | 0.02 | 0.08 | 0.94 | -- |

**Table S10.A.** Analysis: Pre-stimulus FRs for false alarm vs. correct reject in A-rule, Inclusion filter: behavior sessions with >10 FA and CR trials in A-rule.

| Group | *n*<br>(units) | CR<br>trials<br>mean<br>pre-sti<br>m FR<br>(Hz) | ±SD | FA<br>trials<br>mean<br>pre-sti<br>m FR<br>(Hz) | ±SD | WSR<br>Z-value<br>across<br>outcome<br>s | *p*-val | Mean FR<br>difference<br>(Hz):<br>CR-FA<br>(*p*<0.05<br>only) |
| --- | --- | --- | --- | --- | --- | --- | --- | --- |
| **All** |  |  |  |  |  |  |  |  |
| Super. BS | 58 | 3.2 | 3.9 | 3.6 | 3.9 | -1.42 | 0.156 | -- |
| Super. NS | 22 | 15.3 | 10.8 | 15.9 | 10.0 | -0.28 | 0.783 | -- |
| Mid. BS | 78 | 4.2 | 5.2 | 4.3 | 5.2 | -1.32 | 0.189 | -- |
| Mid. NS | 18 | 13.3 | 12.0 | 14.3 | 14.1 | -0.28 | 0.777 | -- |
| Deep BS | 247 | 3.0 | 4.2 | 3.1 | 4.3 | -0.67 | 0.506 | -- |
| Deep NS | 41 | 15.3 | 10.2 | 14.9 | 10.3 | 1.50 | 0.135 | -- |
| **Units with suppressed FRs in A-rule** |  |  |  |  |  |  |  |  |
| Super. BS | 31 | 2.6 | 3.7 | 3.5 | 3.8 | -2.93 | 0.0034 | 0.94 |
| Super. NS | 5 | 9.5 | 7.3 | 13.0 | 8.9 | -2.02 | 0.0431 | 3.49 |
| Mid. BS | 45 | 4.2 | 5.4 | 4.7 | 5.4 | -2.46 | 0.0141 | 0.49 |
| Mid. NS | 11 | 15.6 | 13.7 | 17.8 | 16.2 | -1.16 | 0.2477 | -- |
| Deep BS | 139 | 2.9 | 4.2 | 3.2 | 4.1 | -2.80 | 0.0052 | 0.26 |
| Deep NS | 18 | 12.1 | 7.0 | 12.8 | 7.5 | -0.89 | 0.372 | -- |

**Table S10.B.**
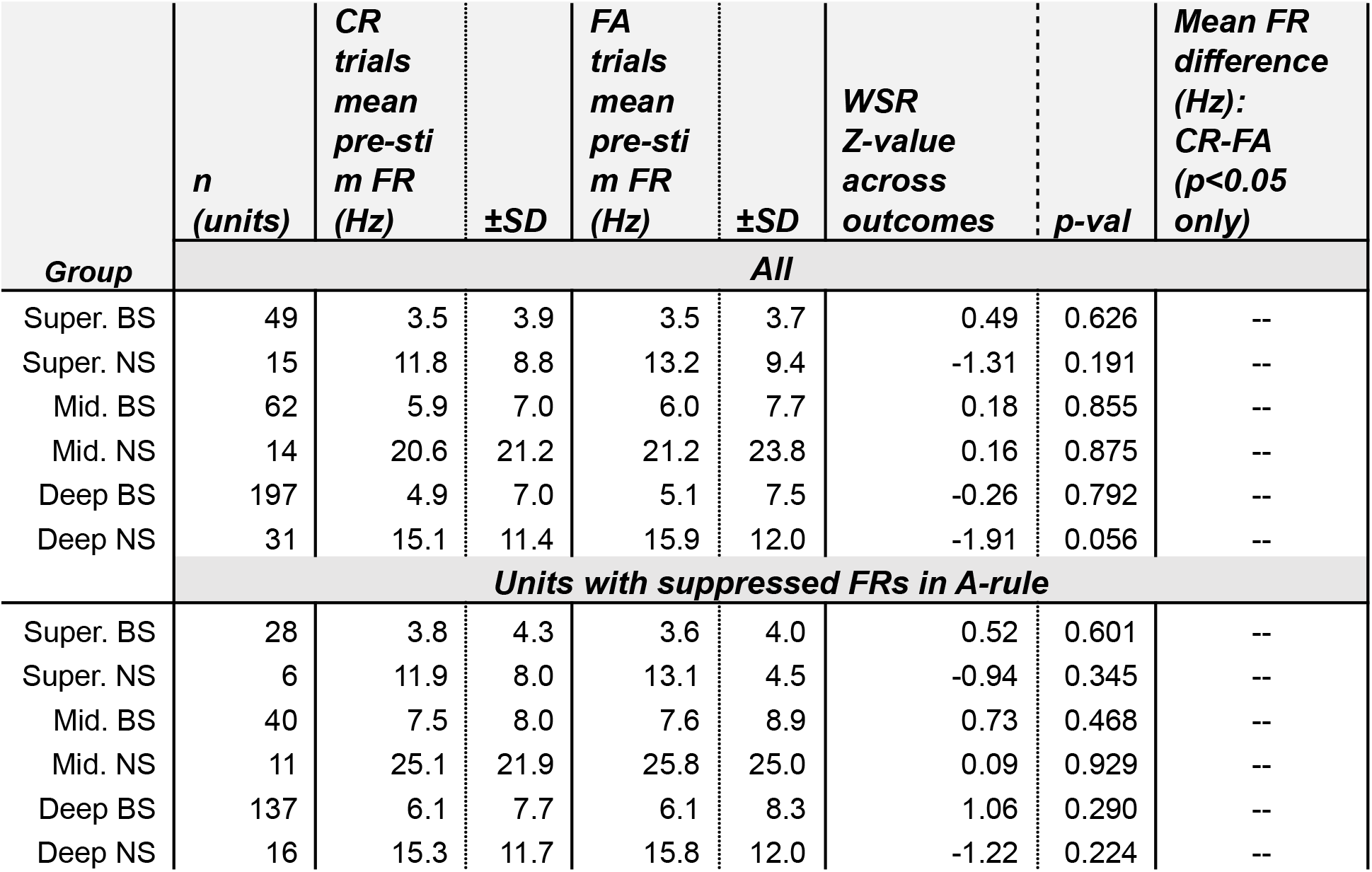
Analysis: Pre-stimulus FRs for false alarm vs. correct reject in V-rule, Inclusion filter: behavior sessions with >10 FA and CR trials in V-rule.

| Group | *n*<br>(units) | CR<br>trials<br>mean<br>pre-sti<br>m FR<br>(Hz) | ±SD | FA<br>trials<br>mean<br>pre-sti<br>m FR<br>(Hz) | ±SD | WSR<br>Z-value<br>across<br>outcomes | p-val | Mean FR<br>difference<br>(Hz):<br>CR-FA<br>($p < 0.05$<br>only) |
| --- | --- | --- | --- | --- | --- | --- | --- | --- |
|  | All |  |  |  |  |  |  |  |
| Super. BS | 49 | 3.5 | 3.9 | 3.5 | 3.7 | 0.49 | 0.626 | -- |
| Super. NS | 15 | 11.8 | 8.8 | 13.2 | 9.4 | -1.31 | 0.191 | -- |
| Mid. BS | 62 | 5.9 | 7.0 | 6.0 | 7.7 | 0.18 | 0.855 | -- |
| Mid. NS | 14 | 20.6 | 21.2 | 21.2 | 23.8 | 0.16 | 0.875 | -- |
| Deep BS | 197 | 4.9 | 7.0 | 5.1 | 7.5 | -0.26 | 0.792 | -- |
| Deep NS | 31 | 15.1 | 11.4 | 15.9 | 12.0 | -1.91 | 0.056 | -- |
| Units with suppressed FRs in A-rule |  |  |  |  |  |  |  |  |
| Super. BS | 28 | 3.8 | 4.3 | 3.6 | 4.0 | 0.52 | 0.601 | -- |
| Super. NS | 6 | 11.9 | 8.0 | 13.1 | 4.5 | -0.94 | 0.345 | -- |
| Mid. BS | 40 | 7.5 | 8.0 | 7.6 | 8.9 | 0.73 | 0.468 | -- |
| Mid. NS | 11 | 25.1 | 21.9 | 25.8 | 25.0 | 0.09 | 0.929 | -- |
| Deep BS | 137 | 6.1 | 7.7 | 6.1 | 8.3 | 1.06 | 0.290 | -- |
| Deep NS | 16 | 15.3 | 11.7 | 15.8 | 12.0 | -1.22 | 0.224 | -- |

